# Titrating gene expression with series of systematically compromised CRISPR guide RNAs

**DOI:** 10.1101/717389

**Authors:** Marco Jost, Daniel A. Santos, Reuben A. Saunders, Max A. Horlbeck, John S. Hawkins, Sonia M. Scaria, Thomas M. Norman, Jeffrey A. Hussmann, Christina R. Liem, Carol A. Gross, Jonathan S. Weissman

**Affiliations:** Department of Cellular and Molecular Pharmacology, University of California, San Francisco, San Francisco, CA, USA; Howard Hughes Medical Institute, University of California, San Francisco, San Francisco, CA, USA; California Institute for Quantitative Biosciences, University of California, San Francisco, San Francisco, CA 94158, USA; Department of Microbiology and Immunology, University of California, San Francisco, San Francisco, CA, USA; Department of Cell and Tissue Biology, University of California, San Francisco, San Francisco, CA, USA

## Abstract

Biological phenotypes arise from the degrees to which genes are expressed, but the lack of tools to precisely control gene expression limits our ability to evaluate the underlying expression-phenotype relationships. Here, we describe a readily implementable approach to titrate expression of human genes using series of systematically compromised sgRNAs and CRISPR interference. We empirically characterize the activities of compromised sgRNAs using large-scale measurements across multiple cell models and derive the rules governing sgRNA activity using deep learning, enabling construction of a compact sgRNA library to titrate expression of ∼2,400 genes involved in central cell biology and a genome-wide *in silico* library. Staging cells along a continuum of gene expression levels combined with rich single-cell RNA-seq readout reveals gene-specific expression-phenotype relationships with expression level-specific responses. Our work provides a general tool to control gene expression, with applications ranging from tuning biochemical pathways to identifying suppressors for diseases of dysregulated gene expression.

---

The complexity of biological processes arises not only from the set of expressed genes but also from quantitative differences in their expression levels. As a classic example, some genes are haploinsufficient and thus are sensitive to a 50% decrease in expression, whereas other genes are permissive to far stronger depletion^1^. Enabled by tools to titrate gene expression levels such as series of promoters or hypomorphic mutants, the underlying expression-phenotype relationships have been explored systematically in yeast^2–4^ and bacteria^5–8^. These efforts have revealed gene- and environment-specific effects of changes in expression levels^4^ and yielded insight into the opposing evolutionary forces that determine gene expression levels including the cost of protein synthesis and the need for robustness against random fluctuations^3, 6, 8^. The availability of equivalent tools in mammalian systems would enable similar efforts to understand these forces in more complex models as well as additional applications. For example, such tools could be used to identify the functionally sufficient levels of gene products, which can serve as targets for rescue by gene therapy or chemical treatment when genes are affected by disease-causing loss-of-function mutations or as targets of inhibition for anti-cancer drugs such that proliferating cancer cells experience toxicity while healthy cells are spared. It is possible to titrate the expression of individual genes in mammalian systems, for example by incorporating microRNA binding sites of varied strength into the 3 -UTR of the endogenous locus^9^. By contrast, functional genomics tools that allow systematic targeting of genes have been primarily optimized for complete knockout or knockdown or strong overexpression and do not afford the required nuanced control over gene expression levels.

The discovery and development of artificial transcription factors, such as TALEs^10^ or the CRISPR-based effectors underlying CRISPR interference (CRISPRi) and activation (CRISPRa)^11^, has brought tools to systematically control gene expression within reach. CRISPR/Cas9 based systems in particular have attracted considerable attention as the targeting to a locus of interest through sequence complementarity to an associated single guide RNA (sgRNA) affords uniquely high programmability^12^. Studies of the targeting mechanisms have established that both activity and binding of Cas9 or its nuclease-dead variants (dCas9) can be modulated by introducing mismatches into the sgRNA targeting region, modifying the sgRNA constant region, and other approaches^12–17^. Here, we report a systematic approach to control dCas9 effector binding through modified sgRNAs as a general method to titrate gene expression in mammalian cells. We describe both an empirically validated compact sgRNA library to titrate the expression of essential genes and a genome-wide *in silico* library derived from deep learning analysis of the empirical data. Rich single-cell RNA-seq phenotypes recorded at different expression levels of essential genes reveal gene-specific expression-phenotype relationships and expression level-dependent cell responses and highlight the utility of such modified sgRNAs in staging cells along a continuum of expression levels.

## Results

### Mismatched sgRNAs mediate diverse intermediate phenotypes

To comprehensively characterize the activities of mismatched sgRNAs in CRISPRi-mediated knockdown, we introduced all 57 singly mismatched variants of a GFP-targeting sgRNA^18^ into GFP^+^ K562 CRISPRi cells and measured GFP levels by flow cytometry (Fig. 1a). Cells harboring mismatched sgRNAs experienced knockdown levels between those of cells with the perfectly matched sgRNA (94%) and cells with a non-targeting control sgRNA (Fig. 1b, S1a-c, Table S1). As expected, sgRNAs with mismatches in the PAM-proximal seed region^12, 13^ had strongly compromised activity. By contrast, sgRNAs with mismatches in the PAM-distal region mediated GFP knockdown to an extent similar to that of the unmodified sgRNA, albeit with substantial variability depending on the type of mismatch (Fig. 1b-c). The distributions of GFP levels with mismatched sgRNAs were largely unimodal, although the distributions were typically broader than with the perfectly matched sgRNA or the control sgRNA (Fig. 1b, S1c). These results suggest that series of mismatched sgRNAs can be used to titrate gene expression at the single-cell level, but that mismatched sgRNA activity is modulated by complex factors.

**Figure 1.**
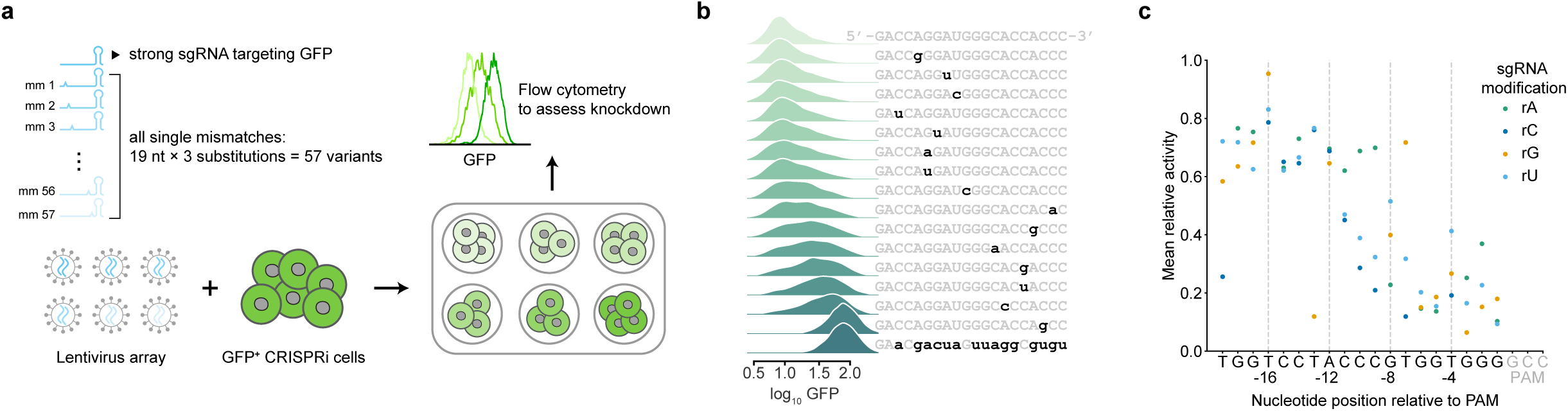
Mismatched sgRNAs titrate GFP expression at the single-cell level. **(a)** Experimental design to test knockdown conferred by all mismatched variants of a GFP-targeting sgRNA. **(b)** Distributions of GFP levels in cells with perfectly matched sgRNA (top), mismatched sgRNAs (middle), and non-targeting control sgRNA (bottom). Sequences of sgRNAs are indicated on the right (without the PAM). **(c)** Relative activities of all mismatched sgRNAs, defined as the ratio of fold-knockdown conferred by a mismatched sgRNA to fold-knockdown conferred by the perfectly matched sgRNA. Relative activities are displayed as the mean of two biological replicates.

### Rules of mismatched sgRNA activity derived from a large-scale screen

We reasoned that we could empirically derive the factors governing the influence of mismatches on sgRNA activity by measuring growth phenotypes imparted by a large number mismatched sgRNAs in a pooled screen. For this purpose, we generated a ∼120,000-element library comprising series of variants for 4,898 sgRNAs targeting 2,499 genes with growth phenotypes in K562 cells^19^. Each individual series, herein referred to as an allelic series, contains the original, perfectly matched sgRNA and 22-23 variants with one or two mismatches (Fig. 2a, Table S2). We then measured CRISPRi growth phenotypes (γ for which a more negative value indicates a stronger growth defect) for each sgRNA in this library in both K562 (chronic myelogenous leukemia) and Jurkat (acute T-cell lymphocytic leukemia) cells using pooled screens^15, 20^ (Fig. 2b, S2a-d, Methods). Growth phenotypes of targeting sgRNAs were well-correlated in biological replicates (Fig. S2a-b, Tables S3-S4, Pearson *r*^2^ [K562] = 0.82; Pearson *r*^2^ [Jurkat] = 0.82) and recapitulated previously reported phenotypes^19^ (Fig. S2c).

**Figure 2.**
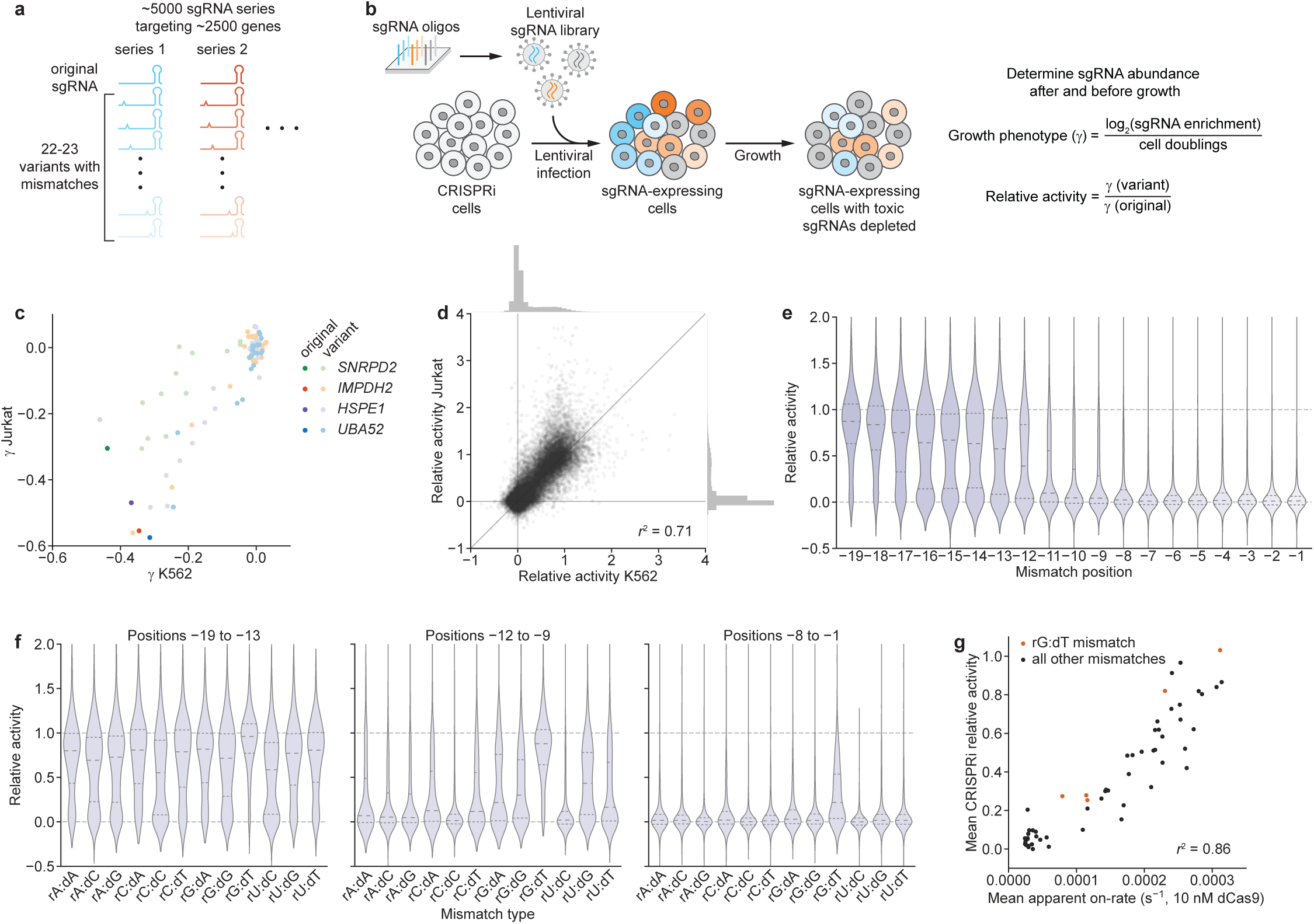
A large-scale CRISPRi screen identifies factors governing mismatched sgRNA activity. **(a)** Design of large-scale mismatched sgRNA library. **(b)** Schematic of pooled CRISPRi screen to determine activities of mismatched-sgRNAs. **(c)** Growth phenotypes (γ) in K562 and Jurkat cells for four sgRNA series, with the perfectly matched sgRNAs shown in darker colors and mismatched sgRNAs shown in corresponding lighter colors. Phenotypes represent the mean of two biological replicates. Differences in absolute phenotypes likely reflect cell type-specific essentiality. **(d)** Comparison of mismatched sgRNA relative activities in K562 and Jurkat cells. Marginal histograms depict distributions of relative activities along the corresponding axes. **(e)** Distribution of mismatched sgRNA relative activities stratified by position of the mismatch. Position –1 is closest to the PAM. **(f)** Distribution of mismatched sgRNA relative activities stratified by type of mismatch, grouped by mismatches located in positions –19 to –13 (PAM-distal region), positions –12 to –9 (intermediate region), and positions –8 to –1 (PAM-proximal/seed region). Division into these regions was based on previous work^12, 13^ and the patterns in Fig. 2e. **(g)** Comparison of mean apparent on-rates measured *in vitro* for mismatched variants of a single sgRNA^22^ and mean relative activities from large-scale screen. Values are compared for identical combinations of mismatch type and mismatch position; mean relative activities were calculated by averaging relative activities for all mismatched sgRNAs with a given combination.

Mismatched sgRNAs mediated a range of phenotypes, spanning from that of the corresponding perfectly matched sgRNA to those of negative control sgRNAs (Fig. 2c). To account for differences in absolute growth phenotypes, we normalized the phenotype of each mismatched sgRNA to that of its corresponding perfectly matched sgRNA (relative activity, Fig. 2b) and filtered for series in which the perfectly matched sgRNA had a strong growth phenotype (Methods). Relative activities measured in K562 and Jurkat cells were well-correlated (Fig. 2d, Pearson *r*^2^ = 0.71), regardless of differences in absolute phenotype of the perfectly matched sgRNAs (Pearson *r*^2^ = 0.74 for |γ[K562] – γ[Jurkat]| > 0.2; Pearson *r*^2^ = 0.70 for |γ[K562] – γ[Jurkat]| < 0.2). We therefore averaged relative activities from both cell lines for further analysis (Methods). Although the majority of mismatched sgRNAs were inactive (Fig. 2d), particularly if they contained two mismatches (Fig. S2e), a substantial fraction exhibited intermediate activity (19,596 sgRNAs with 0.1 < relative activity < 0.9, 25.5% of sgRNAs in series passing filter).

To understand the rules governing the impacts of mismatches on sgRNA activity, we stratified the relative activities of singly-mismatched sgRNAs by properties of the mismatch. As expected, mismatch position was a strong determinant of activity, with mismatches closer to the PAM leading to lower relative activity (Fig. 2e). In agreement with patterns of Cas9 off-target activity^21^, sgRNAs with rG:dT mismatches (A to G mutations in the sgRNA) retained substantial activity even for mismatches close to the PAM (Fig. 2f). Other factors were of lower magnitude and more context-dependent, such as the associations of higher GC content with higher activity for mismatches located 9 or more bases upstream of the PAM (positions –9 to –19), and of mismatch-surrounding G nucleotides with marginally higher activity for mismatches in the intermediate region (Fig. S2f-g). The activities of mismatched sgRNAs thus appear to be determined by general biophysical rules; a premise further supported by the high correlation of relative activities obtained in two different cell lines (Fig. 2d) and the high correlation of mismatched sgRNA activities with previous *in vitro* measurements of dCas9 binding on-rates in the presence of mismatches^22^ (Fig. 2g).

Finally, we evaluated the proportion of sgRNA series that provide access to a range of intermediate CRISPRi growth phenotypes for the targeted gene (relative activity between 0.1 and 0.9). When considering only singly-mismatched sgRNAs, 76.1% of series contain at least 2 sgRNAs with intermediate phenotypes, and that number rises to 86.7% when also including double mismatches (Fig. S2h). As we explored only ∼20% of possible single mismatches and <1% of possible double mismatches, it is likely that intermediate-activity sgRNAs also exist for the remaining series. Altogether, these results suggest that systematically mismatched sgRNAs provide a general method to titrate CRISPRi activity and, consequently, target gene expression.

### Controlling sgRNA activity with modified constant regions

We also explored the orthogonal approach of generating intermediate-activity sgRNAs through modifications to the sgRNA constant region, which is required for binding to Cas9. Although previous work has established that such modifications can lead to increases or decreases in Cas9 activity or have no measurable impact^16, 23–27^, the mutational landscape of the constant region has only been sparsely explored, and largely with the goal of preserving sgRNA activity.

To comprehensively assess the activities of modified sgRNA constant regions, we designed a library of 995 constant region variants comprising all possible single nucleotide substitutions, base pair substitutions, and combinations of these changes (Methods, Table S5) and determined the growth phenotypes for each variant paired with 30 different targeting sequences against 10 essential genes in a pooled screen in K562 cells (Fig. 3a, S3a; Tables S1, S6, S7). We calculated relative activities for each targeting sequence:constant region pair by normalizing its phenotype to that of the targeting sequence paired with the unmodified constant region, identifying 409 constant region variants that on average conferred intermediate activity (0.1-0.9, Fig. 3b). Ten variants selected for individual evaluation also mediated intermediate levels of mRNA knockdown (Fig. S3b). Mapping the activities of constant region variants with single base substitutions onto the structure recapitulated known relationships between constant region structure and function (Fig. 3c). For example, mutation of bases known to mediate contacts^16^ with Cas9 (e.g. the first stem loop or the nexus) generally reduced activity, whereas mutations in regions not contacted by Cas9 (e.g. the hairpin region of stem loop 2) were well-tolerated (Fig. 3c). Notably, several variants carrying mutations in stem loop 2 had consistently increased activities and thus could be useful tools for future applications (Fig. 3b-c).

**Figure 3.**
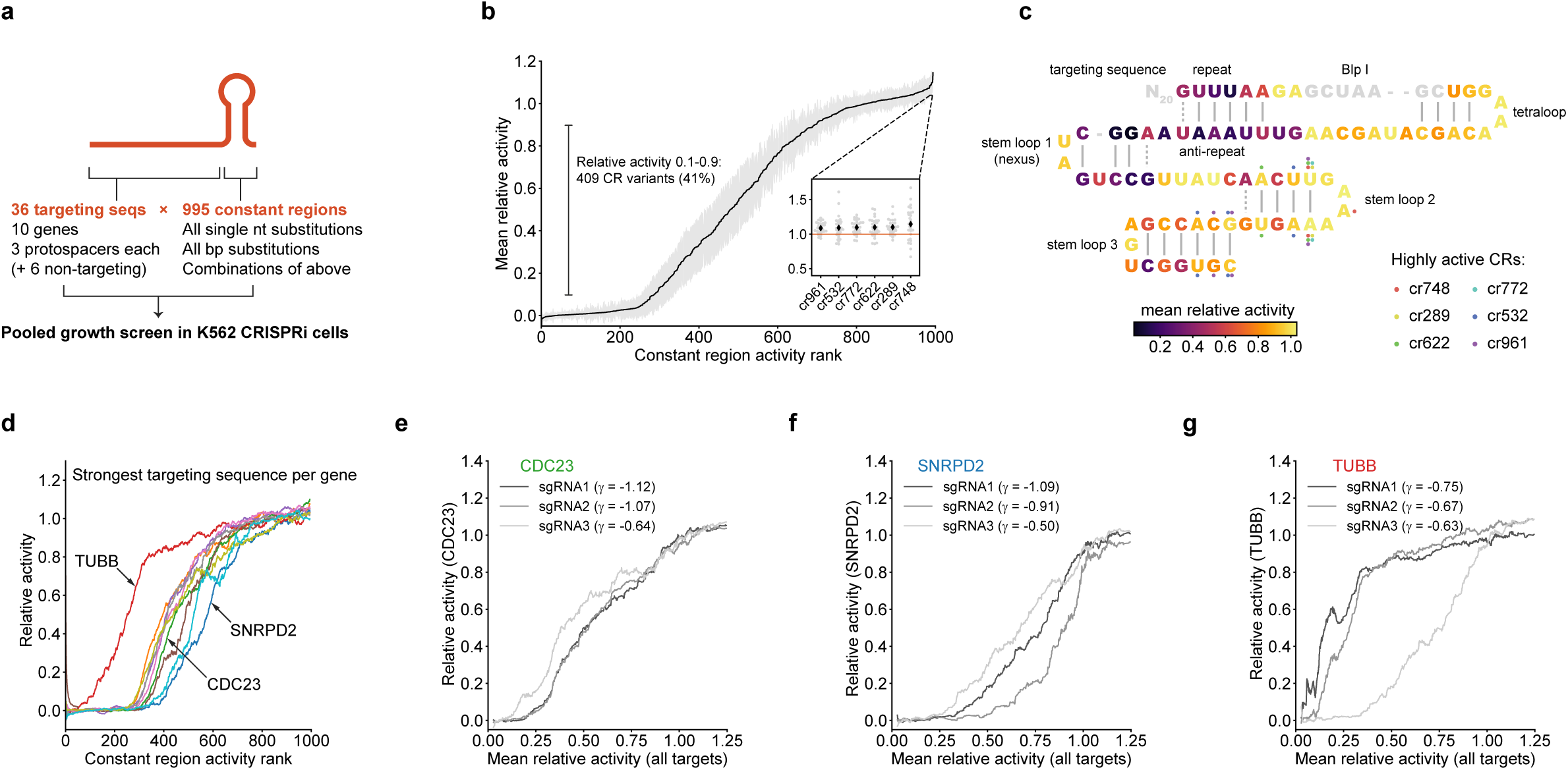
Identification and characterization of intermediate-activity constant regions. **(a)** Design of constant region variant library. **(b)** Mean relative activities of constant region variants, calculated by averaging relative activities for all targeting sequences. Gray margins denote 95% confidence interval. Inset: Focus on 6 constant region variants with higher activity than the original constant region. Black diamonds denote mean relative activity, gray dots relative activities with individual targeting sequences. **(c)** Mapping of constant region variant relative activities onto constant region structure. Each constant region base is colored by the average relative activity of the three single constant region variants carrying a single mutation at that position. Positions mutated in 6 highly active constant regions (inset in panel **b**) are indicated by colored dots. The BlpI site (grey) is used for cloning and was not mutated. **(d)** Constant region activities by targeting sequence, plotted against ranked mean constant region activity. For each gene, the activities with the strongest targeting sequence are shown as rolling means with a window size of 50. **(e-g)** Constant region activities by targeting sequence for all three targeting sequences against the indicated genes. Growth phenotypes (γ) of each targeting sequence paired with the unmodified constant region are indicated in the legend.

Evaluating the relative activities of constant region variants across different targeting sequences revealed consistent rank ordering but substantial variation in the actual values (Fig. 3d, S3c). For example, a targeting sequence against *TUBB* retained high activity with ∼100 constant region variants that otherwise abolished activity for other targeting sequences, whereas a targeting sequence against *SNRPD2* lost activity with ∼50 variants that otherwise conferred intermediate activity (Fig. 3d). In some but not all (Fig. 3e) cases, this heterogeneity extended to different targeting sequences against the same gene, both at the level of growth phenotype (Fig. 3f-g, S3d-e) and mRNA knockdown (Fig. S3b). These differences between targeting sequences could be a consequence of specific targeting sequence:constant region structural interactions or of differences in basal sgRNA expression levels such that lowly expressed sgRNAs are more susceptible to constant region modifications. Thus, although modified constant regions can be used to titrate gene expression, the activity of a given constant region variant for a given targeting sequence is difficult to predict. We therefore focused on sgRNAs with mismatches in the targeting region for the remainder of our work, given that the activities of these sgRNAs were governed by biophysical principles, which should be more predictable.

### A neural network predicts mismatched sgRNA activities with high accuracy

We next sought to leverage our large-scale data set of mismatched sgRNA activities to learn the underlying rules in a principled manner and to enable predictions of intermediate-activity sgRNAs against other genes. We reasoned that a convolutional neural network (CNN) would be well-suited to uncovering these rules due to the ability of CNNs to learn complex global and local dependencies on spatially-ordered features such as nucleotide sequences^28^, including factors governing guide RNA activity in orthogonal CRISPR systems^29, 30^.

To develop a CNN model capable of predicting mismatched sgRNA activities, we constructed a model consisting of two convolution steps, a pooling step, and a 3-layer fully connected neural network (Fig. 4a, S4a). As inputs, the model received sgRNA relative activities paired with nucleotide sequences represented by binarized 3D arrays denoting the genomic sequence of the target and the associated sgRNA mismatch (Fig. 4a). After optimizing hyperparameters using a randomized grid search, we trained 20 independent, equivalently initialized models on the same set of randomly selected sgRNAs (80% of all series) for 8 epochs, which minimized loss without extensive over-fitting (Fig. S4b). Predicted and measured sgRNA relative activities for the validation sgRNA set (i.e. the remaining 20% of series that were not used to train the model) were well-correlated (Pearson *r*^2^ = 0.65), with mean predictions of the 20-model ensemble outperforming all individual models (Fig. 4b, S4c). The distribution of correlation coefficients for individual sgRNA series was unimodal with Pearson *r* values in the 25^th^-75^th^ percentile ranging from 0.77 to 0.93, indicating that the model performed comparably well for most series (Fig. 4c). Model accuracy varied by mismatch position and type, with the highest accuracies corresponding to mismatches in the PAM-proximal seed region (Fig S4d-e). Despite the fact that the model was trained on relative growth phenotypes, it also accurately predicted relative fluorescence values measured in the GFP experiment (Fig. 4d), further supporting the hypothesis that relative growth phenotypes report on biophysical attributes of specific sgRNA:DNA interactions.

**Figure 4.**
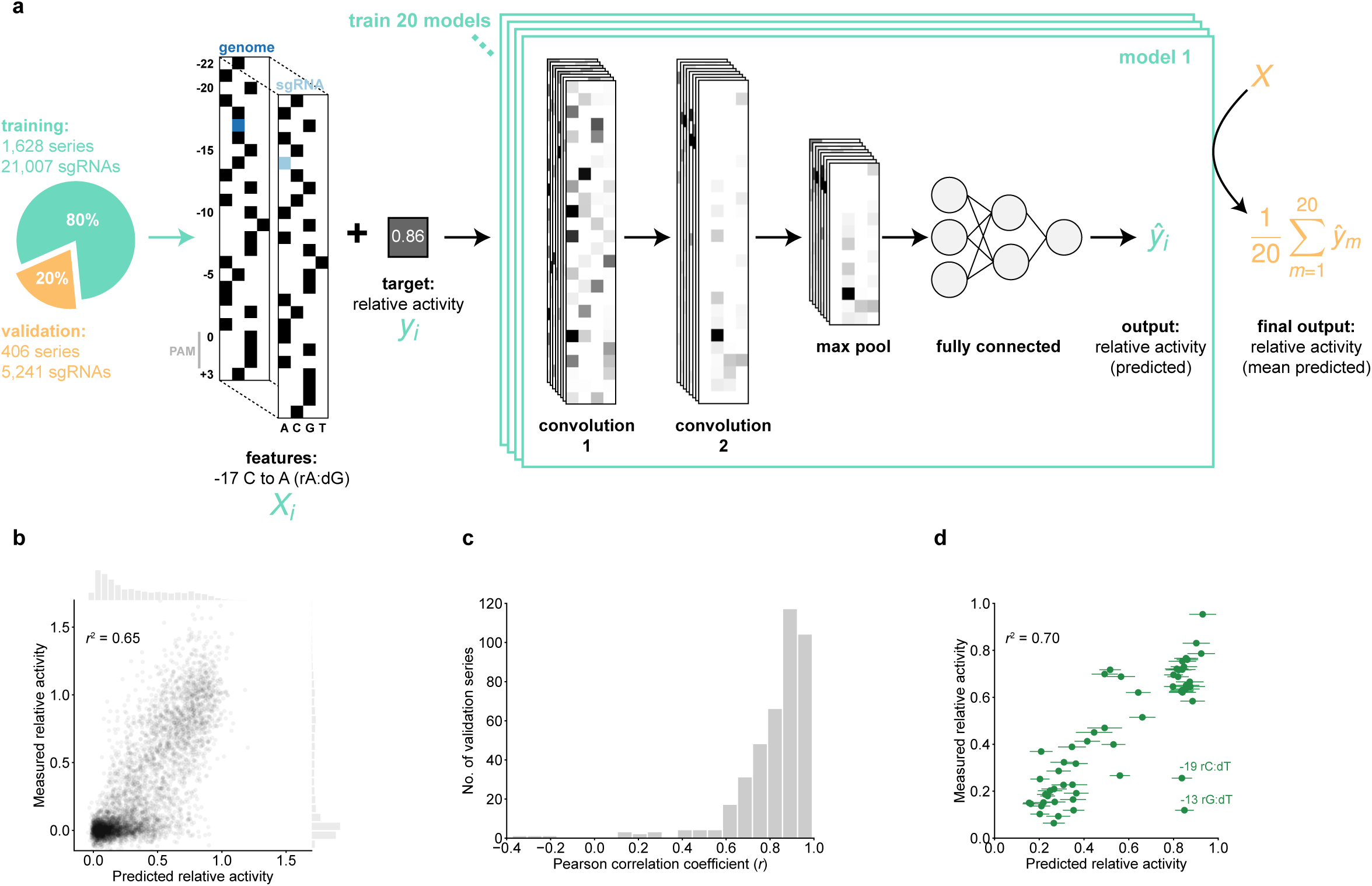
Neural network predictions of sgRNA activity. **(a)** Schematic of a singly-mismatched sgRNA feature array (*X_i_*) and the convolutional neural network architecture trained on pairs of such arrays and their corresponding relative activities (*y_i_*). Black squares in *X_i_* represent the value 1 (the presence of a base at the indicated position); white represents 0. The mean prediction from 20 independently trained models was used to assign a final prediction (ŷ) to each sgRNA in the hold-out validation set (orange). **(b)** Comparison of measured relative growth phenotypes from the large-scale screen and predicted activities assigned by the neural network. Marginal histograms show distributions of relative activities along the corresponding axes**. (c)** Distribution of Pearson *r* values (predicted vs. measured relative activity) for each sgRNA series in the validation set. **(d)** Comparison of measured relative activity (i.e. relative knockdown) in the GFP experiment and predicted relative sgRNA activity. Two outliers with lower-than-predicted activity are annotated with their respective mismatch position and type. Predictions are shown as mean ± S.D. from the 20-model ensemble.

To derive intermediate-activity sgRNAs for all human genes, we used the CNN ensemble to predict relative activities for all 57 singly-mismatched sgRNAs for the top 5 sgRNAs against each gene in the hCRISPRi-v2.1 library^19^ (Table S8). Based on the accuracy of predictions for the validation set, we estimate that for any given gene, sampling 5 sgRNAs with predicted intermediate relative activity (0.1-0.9) will yield at least one sgRNA in that activity range over 90% of the time (Fig. S4f-i). This resource should therefore enable titrating the expression of any gene(s) of interest.

Finally, we sought to further understand the features of mismatched sgRNAs that contribute most to their activity. As the contributions of individual features in a deep learning model are difficult to assess directly, we also trained an elastic net linear regression model on the same data using a curated set of features. This linear model explained less variance in relative activities than the CNN model (*r*^2^ = 0.52, Fig. S5a-b), implying that our feature set was incomplete and/or sgRNA activity is partly determined by non-linear combinations of features; nonetheless, the relative activities predicted by the different models were well-correlated (*r*^2^ = 0.74, Fig. S5c). Consistent with our earlier observations, mismatch position and type were assigned the largest absolute weights in the model, although other features such as GC content in the sgRNA and the identities of flanking bases up to 3 nucleotides away from the mismatch were heavily weighted as well (Fig. S5d-e). For any given position, the type of mismatch contributed differentially to the prediction, with the largest variation between types occurring in the intermediate region of the targeting sequence (Fig. S5f). Taken together, these data demonstrate that the activities of mismatch-containing sgRNAs are determined by multiple factors which can be captured using supervised machine learning approaches.

### A compact mismatched sgRNA library conferring intermediate growth phenotypes

We next set out to design a more compact version of our large-scale library to titrate essential genes with a limited number of sgRNAs. We selected 2,405 genes which we had found to be essential for robust growth in K562 cells in our large-scale screen, divided the relative activity space into six bins, and attempted to select mismatched variants from each of the center four bins (relative activities 0.1-0.9) for two sgRNA series targeting each gene. If a bin did not contain a previously measured sgRNA, we selected one from the CNN model ensemble predictions (Fig. 5a), filtered to exclude sgRNAs with off-target binding potential. For each gene, 2 perfectly matched and 8 mismatched sgRNAs were selected, with approximately 32% of mismatched sgRNAs imputed from the CNN model (Fig. S6a-c, Table S9).

**Figure 5.**
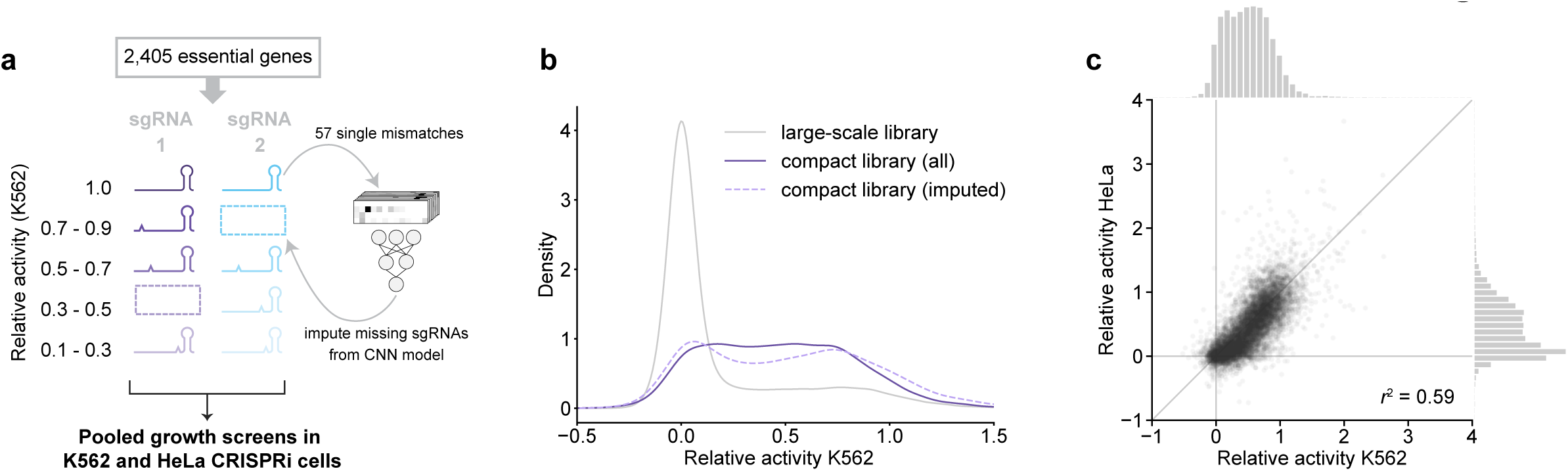
Compact mismatched sgRNA library targeting essential genes. **(a)** Design of library. For activity bins lacking a previously measured sgRNA, novel mismatched sgRNAs were included according to predicted activity. **(b)** Distribution of relative activities from the large-scale library (gray) and the compact library (purple) in K562 cells. **(c)** Comparison of relative activities of mismatched sgRNAs in HeLa and K562 cells. Marginal histograms show the distributions of relative activities along the corresponding axes.

We evaluated the relative activities of sgRNAs in the compact library using pooled CRISPRi growth screens in K562 and HeLa (cervical carcinoma) cells (Tables S10, S11). Growth phenotypes were well-correlated in biological replicates from samples harvested at different time points after t_0_ in both cell lines (Fig. S6d-f). The CNN model predicted imputed sgRNA activities with lower accuracy than the large-scale validation (Fig. S6g), although we note that imputed sgRNAs were highly enriched in PAM-distal mutations which are associated with higher model errors (Fig. S6b, S4e). Whereas the majority of mismatched sgRNAs in the large-scale screen had little to no activity, relative activities in the compact library were evenly distributed, ranging from inactive to full activity (Fig. 5b). Relative sgRNA activities were also well-correlated between K562 and HeLa cells (*r*^2^ = 0.58, Fig. 5c), suggesting that our library provides access to intermediate phenotypes for this core set of genes in multiple cell types.

### Exploring expression-phenotype relationships with sgRNA series

Finally, we sought to use intermediate-activity sgRNAs to explore relationships between expression levels of various genes and the resulting cellular phenotypes. To simultaneously measure gene expression levels and obtain rich phenotypes for a variety of sgRNA series, we used Perturb-seq, an experimental strategy that enables matched capture of the transcriptome and the identity of an expressed sgRNA for each individual cell in pools of cells^27, 31–33^ (Fig. S7a). We chose 25 essential genes involved in diverse cell biological processes (Table S12), targeting each with a perfectly matched sgRNA and 4-5 variants with intermediate growth phenotypes (138 sgRNAs total including 10 non-targeting controls, Table S1). We then subjected pooled K562 CRISPRi cells expressing these sgRNAs from a modified CROP-seq vector^33, 34^ to single-cell RNA-seq (scRNA-seq), using the co-expressed sgRNA barcodes to assign unique sgRNA identities to ∼19,600 cells (median 122 cells per sgRNA, Fig. S7b-c). In addition to the single-cell transcriptomes, we measured bulk growth phenotypes conferred by sgRNAs in these cells. These growth phenotypes were well-correlated with those from the large-scale screen and were used to assign sgRNA relative activities for further analysis (Methods, Fig. S7d-e, Table S13, S14).

We first used the scRNA-seq data to assess the expression of the gene targeted by each sgRNA series. To account for cell-to-cell variability in transcript capture efficiency, we quantified target gene UMIs as a fraction of total UMIs in a given cell (Fig. S8), although analyzing raw UMI counts yielded similar results (Fig. S9). Approximately half of the genes we targeted were highly expressed (median >10 UMIs per cell), allowing us to directly measure target gene expression levels on the single-cell level (Fig. 6a, S8). These distributions are largely unimodal, with medians shifting downwards with increasing sgRNA activity (Fig. 6a). For some of these genes, however, two populations with different knockdown levels are apparent (Fig. 6a, S8a). These populations are present both with intermediate-activity sgRNAs and the perfectly matched sgRNAs, suggesting that they are not a consequence of limited knockdown penetrance for intermediate-activity sgRNAs. Owing to the limited capture efficiency of scRNA-seq, for genes with intermediate to low expression such as *CAD* and *COX11* we typically observed 0-4 UMIs per cell, rendering the quantification of single-cell expression levels more difficult. We nonetheless observe a shift of the distribution to lower UMI numbers with increasing sgRNA activity (Fig. S8a, S9) as well as a decrease in mean expression levels when averaging expression across all cells with the same sgRNA (Fig. S8b).

**Figure 6.**
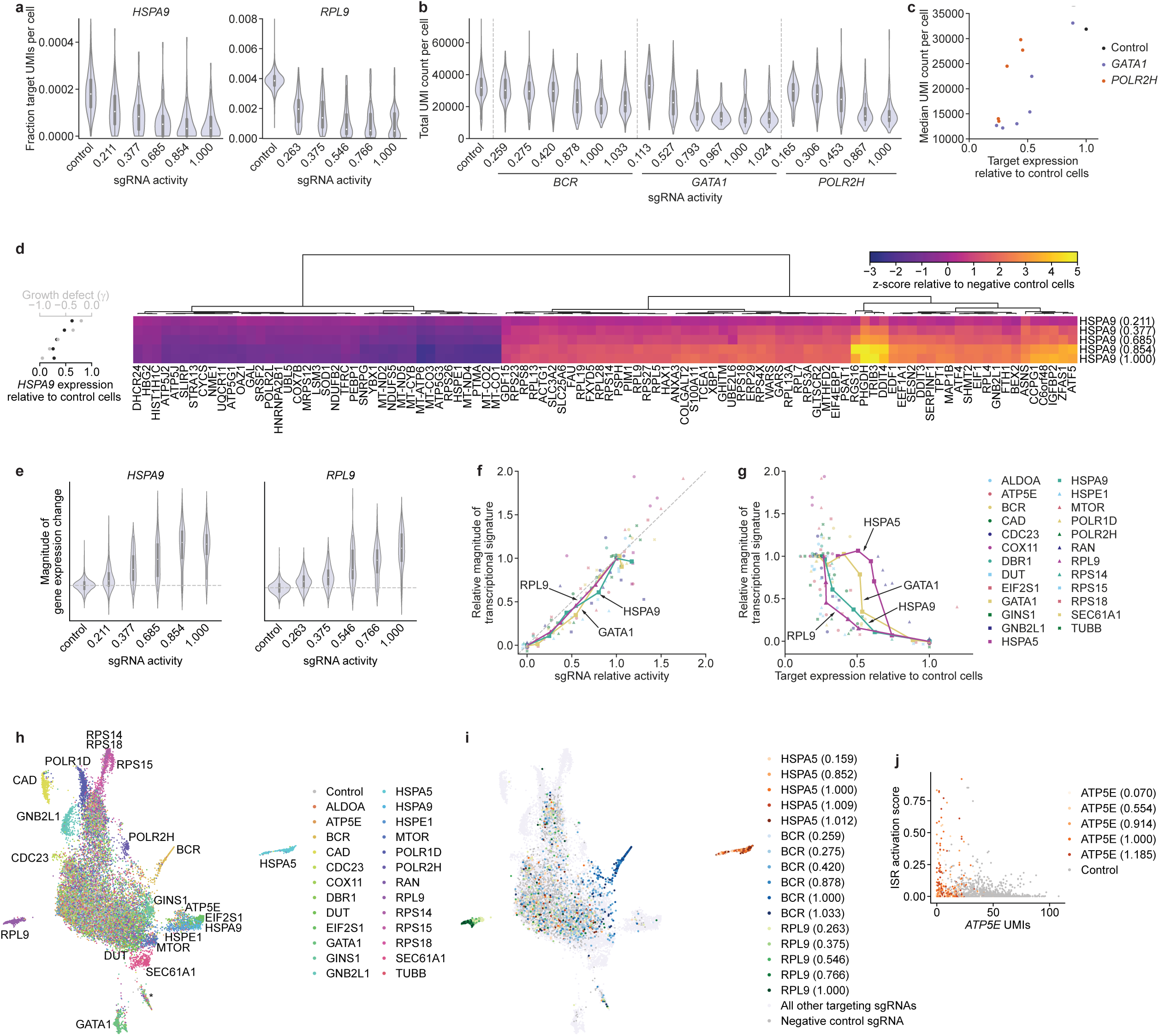
Rich phenotyping of cells with intermediate-activity sgRNAs by Perturb-seq. **(a)** Distributions of *HSPA9* and *RPL9* expression in cells with indicated perturbations. Expression is quantified as target gene UMI count normalized to total UMI count per cell. sgRNA activity is calculated using relative γ measurements from the Perturb-seq cell pool after 5 days of growth. **(b)** Distributions of total UMI counts in cells with indicated perturbations. **(c)** Comparison of median UMI count per cell and target gene expression in cells with *GATA1*- or *POLR2H*-targeting sgRNAs. **(d)** Right: Expression profiles of 100 genes in populations with *HSPA9*-targeting sgRNAs. Genes were selected by lowest FDR-corrected *p*-values in cells with the strongest sgRNA from a two-sided Kolmogorov-Smirnov test (Methods). Expression is quantified as z-score relative to population of cells with non-targeting sgRNAs. Left: Growth phenotype and knockdown for each sgRNA. **(e)** Distribution of gene expression changes in populations with indicated sgRNAs. Magnitude of gene expression change is calculated as sum of z-scores of genes differentially expressed in the series (FDR-corrected *p* < 0.05 with any sgRNA in the series, two-sided Kolmogorov-Smirnov test, Methods), with z-scores of individual genes signed by the direction of change in cells with the perfectly matched sgRNA. Distribution for negative control sgRNAs is centered around 0 (dashed line). **(f)** Comparison of relative growth phenotype and magnitude of gene expression change for all individual sgRNAs. Growth phenotype and magnitude of gene expression change are normalized in each series to those of the sgRNA with the strongest knockdown. Individual series highlighted as indicated. **(g)** Comparison of magnitude of gene expression and target gene knockdown, as in **f**. **(h)** UMAP projection of all single cells with assigned sgRNA identity in the experiment, colored by targeted gene. Clusters clearly assignable to a genetic perturbation are labeled. Cluster labeled * contains a small number of cells with residual stress response activation and could represent apoptotic cells. Note that ∼5% cells appear to have confidently but wrongly assigned sgRNA identities, as evident within the cluster of cells with *HSPA5* knockdown (Methods). Given the strong trends in the other results, we concluded that such misassignment did not substantially affect our ability to identify trends within cell populations and in the future may be avoided by approaches to directly capture the expressed sgRNA^34^. **(i)** UMAP projection, as in **h**, with selected series colored by sgRNA activity. **(k)** Comparison of extent of ISR activation to *ATP5E* UMI count in cells with knockdown of *ATP5E* or control cells.

Titration is also apparent at the level of the transcriptional responses, which provides a robust single-cell measurement of the phenotype induced by depletion of the targeted gene. In the simplest cases, knockdown led to substantial reductions in cellular UMI counts, consistent with large-scale inhibition of mRNA transcription (Fig. 6b, Fig. S10a). Examples include *GATA1*, a central myeloid lineage transcription factor, *POLR2H*, a core subunit of RNA polymerase II (as well as RNA polymerases I and III), or to a lesser extent *BCR*, which is fused to the driver oncogene *ABL1* in K562 cells^35, 36^. Notably, this effect correlates linearly with growth phenotype for intermediate activity sgRNAs (Fig. 6b, Fig. S10b) but exhibits non-linear relationships with target gene knockdown at least in the cases of *GATA1* and *POLR2H* (Fig. 6c, S10b, *BCR* levels are difficult to quantify accurately). Both relationships appear to be sigmoidal but with different thresholds: whereas cellular UMI counts drop rapidly once *GATA1* mRNA levels are reduced by 50%, a larger reduction of *POLR2H* mRNA levels is required to achieve a similarly sized effect. Knockdown of most other targeted genes did not perturb total UMI counts to the same extent (Fig. S10a) but resulted in other transcriptional responses. Knockdown of *CAD*, for example, triggered cell cycle stalling during S-phase, as had been observed previously^27^, with a higher frequency of stalling with increasing sgRNA activity (Fig. S10c). By contrast, knockdown of *HSPA9*, the mitochondrial Hsp70 isoform, induced the expected transcriptional signature corresponding to activation of the integrated stress response (ISR) including upregulation of *DDIT3* (CHOP), *DDIT4*, *ATF5*, and *ASNS*^27, 37^. The magnitude of this transcriptional signature increased with increasing sgRNA activity on both the bulk population (Fig. 6d) and single-cell levels (Fig. 6e), although populations with intermediate-activity sgRNAs had larger cell-to-cell variation in the magnitudes of transcriptional responses. Similarly, the transcriptional responses to knockdown of other genes (Fig. S10d) scaled with sgRNA activity and exhibited larger variance for intermediate-activity sgRNAs (Fig. 6e).

We next explored expression-phenotype relationships in these data. Within each series, two major metrics of phenotype, bulk population growth phenotype and transcriptional response, appear to be well-correlated, despite substantial differences in the absolute magnitudes of the transcriptional responses with different series (Fig. 6f, S10d-f). By contrast, the relationships between either metric of phenotype and target gene expression are strongly gene-specific (Fig. 6g, Fig. S10g-i). For *HSPA5* and *GATA1*, for example, a comparably small reduction in mRNA levels (∼50%) was sufficient to induce a near-maximal transcriptional response and growth defect, whereas for most other genes a larger reduction was required. These results prompt the hypothesis that K562 cells are intolerant to moderate decreases in expression of *GATA1* and *HSPA5*, with sharp transitions from growth to death once expression levels drop below a threshold. More broadly, these results highlight the utility of titrating gene expression to systematically map expression-phenotype relationships and quantitatively define gene expression sufficiency.

### Following single-cell trajectories along a continuum of gene expression levels

To gain further insight into the diversity of transcriptional responses induced by depletion of essential genes, we compared the transcriptional profiles of all perturbations. Clustering perturbations according to the similarity (Pearson correlation) of their bulk transcriptomes revealed multiple groups segregated by biological function, including a cluster of ribosomal proteins and *POLR1D*, a subunit of the rRNA-transcribing RNA polymerase I (and of RNA polymerase III), and a cluster of perturbations that activate the integrated stress response *HSPA9, HSPE1*, and *EIF2S1*/eIF2α) (Fig. S11a). To further visualize the space of α transcriptional states, we performed dimensionality reduction on the single-cell transcriptomes using UMAP^38^. The resulting projection recapitulates the clustering, as indicated for example by the close proximity of cells with perturbations of *HSPA9*, *HSPE1*, and *EIF2S1* (Fig. 6h). Within individual series, cells project further outward in UMAP space with increasing sgRNA activity, further highlighting that target gene expression levels are titrated on the single cell level (Fig. 6i).

Closer examination of the UMAP projection revealed more granular structure, including the grouping of a subset of cells with knockdown of *ATP5E*, a subunit of ATP synthase, with cells with ISR-activating perturbations (Fig. 6h). This subset of cells indeed exhibited classical features of ISR activation (Fig. S11b). The frequency of ISR activation increased with lower *ATP5E* mRNA levels, but even at the lowest levels some cells did not exhibit ISR activation (Fig. 6j, S11b). These results suggest that depletion of ATP synthase under these conditions predisposes cells to activate the ISR, perhaps by exacerbating transient phases of mitochondrial stress, in a manner that is proportional to ATP synthase levels. More broadly, these results highlight the utility of titrating gene expression in probing cell biological phenotypes, especially in combination with rich phenotyping methods such as scRNA-seq.

## Discussion

Here we describe the development of allelic series of compromised sgRNAs, with each series enabling the titration of the expression of a given gene in human cells. These series, either individually or as a pool, have a broad range of applications across basic and biomedical research. We highlight the utility of the approach in extracting rich phenotypes by single-cell RNA-seq along a continuum of gene expression levels, which enabled mapping of expression levels to various phenotypes and identification of expression level-dependent cell fates.

Our approach builds on *in vitro* work describing the biophysical principles by which modifications to the sgRNA modulate (d)Cas9 binding on-rates and activity^13, 22, 39–41^. In cells, modifications to the sgRNA constant region were affected by specific interactions with targeting sequences, rendering sgRNA activities difficult to predict. By contrast, the effects of mismatches on sgRNA activity followed more readily discernable biophysical principles, enabling us to apply machine learning approaches to derive the underlying rules and predict series for arbitrary sgRNAs. The resulting genome-wide *in silico* library (Table S8) enables titration of any expressed gene of interest. We also describe a compact (25,000-element) library that enables titration of ∼2,400 essential genes (Table S9), with potential applications for example in focused screens for sensitization to chemical or genetic perturbations. Given that target gene expression levels are largely unimodally distributed in cell populations harboring sgRNA series, these sgRNAs can be combined with both single-cell or bulk population readouts. Thus, complex phenotypes as a function of gene expression levels can be recorded by a variety of techniques tailored to the particular question, such as Perturb-seq or related techniques, microscopy, bulk metabolomics or proteomics, or targeted cell biological assays, providing substantial experimental flexibility.

These sgRNA series now enable mapping expression-to-phenotype curves directly in mammalian systems, with implications for example for evolutionary biology and biomedical research. Indeed, using sgRNA series to titrate essential gene expression, we found gene-specific expression-phenotype relationships: although all genes had a threshold expression level below which cell viability dropped rapidly, the relative locations of these thresholds varied across genes, with K562 cells being particularly sensitive to depletion of *GATA1* and *HSPA5*. This variability in threshold location suggests different buffering capacities for different genes, in line with previous findings in yeast^4^, but the logic by which these buffering capacities are determined in mammalian systems remains unclear. More comprehensive efforts to generate such dose-response curves and determine the extents to which gene expression is buffered across cell models would allow for identification of patterns for different gene sets and biological processes and thereby begin to reveal the underlying principles that have shaped gene expression levels. Analogous efforts to map such dose-response curves in cancer cell types could identify specific vulnerabilities as targets for therapeutics and, vice versa, mapping these curves for cancer driver genes or genes underlying specific diseases could enable defining the corresponding therapeutic windows, i.e. the required extents of inhibition or restoration, as goals for drug development.

Our intermediate-activity sgRNAs also provide access to a diversity of cell states including loss-of-function phenotypes that otherwise may be obscured by cell death or neomorphic behavior. Thus, our approach enables positioning cells at states of interest, for example to record chemical-gene or gene-gene interactions, or near phenotypic transitions to characterize the transcriptional trajectories. These sgRNA series will also facilitate recapitulating gene expression levels of disease-relevant states such as haploinsufficiency or partial loss-of-function diseases, enabling systematic efforts to identify suppressors or modifiers as potential therapeutic targets, or modeling quantitative trait loci associated with multigenic traits in conjunction with rich phenotyping to systematically identify the mechanisms by which they interact and contribute to such traits. Finally, sgRNA allelic series can be equivalently used to titrate dCas9 occupancy and activity in other applications such as CRISPRa or dCas9-based epigenetic modifiers.

More generally, our allelic series approach now provides a tool to systematically titrate gene expression and evaluate dose-response relationships in mammalian systems. This resource should be equally enabling to systematic large-scale efforts and detailed single-gene investigations in basic cell biology, drug development, and functional genomics.

## Online methods

### Reagents and cell lines

K562 and Jurkat cells were grown in RPMI 1640 medium (Gibco) with 25 mM HEPES, 2 mM L-glutamine, 2 g/L NaHCO_3_ supplemented with 10% (v/v) standard fetal bovine serum (FBS, HyClone or VWR), 100 units/mL penicillin, 100 µg/mL streptomycin, and 2 mM L-glutamine (Gibco). HEK293T and HeLa cells were grown in Dulbecco’s modified eagle medium (DMEM, Gibco) with 25 mM D-glucose, 3.7 g/L NaHCO_3_, 4 mM L-glutamine and supplemented with with 10% (v/v) FBS, 100 units/mL penicillin, 100 µg/mL streptomycin, and 2 mM L-glutamine. K562 and HeLa cells are derived from female patients. Jurkat cells are derived from a male patient. HEK293T are derived from a female fetus. K562 and HeLa CRISPRi cell lines were previously published^15, 18^. Jurkat CRISPRi cells (Clone NH7) were obtained from the Berkeley Cell Culture Facility. All cell lines were grown at 37 °C in the presence of 5% CO_2_. K562, Jurkat, and HEK293T cell lines were periodically tested for Mycoplasma contamination using the MycoAlert Plus Mycoplasma detection kit (Lonza). HeLa CRISPRi cells had previously been tested for Mycoplasma contamination but were not explicitly tested as part of this work.

### DNA transfections and virus production

Lentivirus was generated by transfecting HEK239T cells with four packaging plasmids (for expression of VSV-G, Gag/Pol, Rev, and Tat, respectively) as well as the transfer plasmid using TransIT®-LT1 Transfection Reagent (Mirus Bio). Viral supernatant was harvested two days after transfection and filtered through 0.44 µm PVDF filters and/or frozen prior to transduction.

### Cloning of individual sgRNAs

Individual perfectly matched or mismatched sgRNAs were cloned essentially as described previously^15^. Briefly, two complementary oligonucleotides (Integrated DNA Technologies), containing the targeting region as well as overhangs matching those left by restriction digest of the backbone with BstXI and BlpI, were annealed and ligated into an sgRNA expression vector digested with BstXI (NEB or Thermo Fisher Scientific) and BlpI (NEB) or Bpu1102I (Thermo Fisher Scientific). The ligation product was transformed into Stellar™ chemically competent *E. coli* cells (Takara Bio) and plasmid was prepared following standard protocols.

### Individual evaluation of sgRNA phenotypes for GFP knockdown

For individual evaluation of GFP knockdown phenotypes, sgRNAs were individually cloned as described above, ligated into a version of pU6-sgCXCR4-2 (marked with a puromycin resistance cassette and mCherry, Addgene #46917)^18^, modified to include a BlpI site. Sequences used for individual evaluation are listed in Table S1. The sgRNA expression vectors were individually packaged into lentivirus and transduced into GFP^+^ K562 CRISPRi cells^18^ at MOI < 1 (15 – 40% infected cells) by centrifugation at 1000 × *g* and 33 °C for 0.5-2 h. GFP levels were recorded 10 d after transduction by flow cytometry using a FACSCelesta flow cytometer (BD Biosciences), gating for sgRNA-expressing cells (mCherry^+^). Experiments were performed in duplicate from the transduction step. Relative activities were defined as the fold-knockdown of each mismatched variant (GFP_sgRNA_[_non-targeting_] / GFP_sgRNA_[_variant_]) divided by the fold-knockdown of the perfectly-matched sgRNA. The background fluorescence of a GFP^−^ strain was subtracted from all GFP values prior to other calculations. Data were analyzed in Python 2.7 using the FlowCytometryTools package (v0.5.0). The distributions of GFP values in Fig. 1B were plotted following the example in https://seaborn.pydata.org/examples/kde_ridgeplot.

### Design of large-scale mismatched sgRNA library

To generate the list of targeting sgRNAs for the large-scale mismatched sgRNA library, hit genes from a growth screen performed in K562 cells with the CRISPRi v2 library^19^ were selected by calculating a discriminant score (phenotype z-score × –log_10_(Mann-Whitney *P*)). Discriminant scores for negative control genes (randomly sampled groups of 10 non-targeting sgRNAs) were calculated as well, and hit genes were selected above a threshold such that 5% of the hits would be negative control genes (i.e. an estimated empirical 5% FDR). This procedure resulted in the selection of 2477 genes. Of these genes, 28 genes for which the second strongest sgRNA by absolute value had a positive growth phenotype were filtered out as these were likely to be scored as hits solely due to a single sgRNA. For the remaining 2,449 genes, the two sgRNAs with the strongest growth phenotype were selected, for a total of 4,898 perfectly matched sgRNAs.

For each of these sgRNAs, a set of 23 variant sgRNAs with mismatches was designed: 5 with a single randomly chosen mismatch within 7 bases of the PAM, 5 with a single randomly chosen mismatch 8-12 bases from the PAM, and 3 with a single randomly chosen mismatch 13-19 bases from the PAM (the first base of the targeting region was never selected for this purpose as it is an invariant G in all sgRNAs to enable transcription from the U6 promoter). The remaining 10 variants had 2 randomly chosen mismatches selected from positions –1 to –19.

To assess the off-target potential of mismatched sgRNAs, we extended our previous strategy to estimate sgRNA off-target effects^15, 19^. Briefly, for each target in the genome, a FASTQ entry was created for the 23 bases of the target including the PAM, with the accompanying empirical Phred score indicating an estimate of the anticipated importance of a mismatch in that base position. Bowtie (http://bowtie-bio.sourceforge.net)42 was then used to align each designed sgRNA back to the genome, parameterized so that sgRNAs were considered to mutually align if and only if: a) no more than 3 mismatches existed in the PAM-proximal 12 bases and the PAM, b) the summed Phred score of all mismatched positions across the 23 bases was less than a threshold. This alignment was done iteratively with decreasing thresholds, and any sgRNAs which aligned successfully to no other site in the genome at a particular threshold were then deemed to have a specificity at said threshold. The compiled sgRNA sequences were then filtered for sgRNAs containing BstXI, BlpI, and SbfI sites, which are used during library cloning and sequencing library preparation, and 2,500 negative controls (randomly generated to match the base composition of our hCRISPRi-v2 library) were added. Sequences of sgRNAs and descriptions of mismatches are listed in Table S2.

### Pooled cloning of mismatched sgRNA libraries

Pooled sgRNA libraries were cloned largely as described previously^15, 20, 43^. Briefly, oligonucleotide pools containing the desired elements with flanking restriction sites and PCR adapters were obtained from Agilent Technologies. The oligonucleotide pools were amplified by 15 cycles of PCR using Phusion polymerase (NEB). The PCR product was digested with BstXI (Thermo Fisher Scientific) and Bpu1102I (Thermo Fisher Scientific), purified, and ligated into BstXI/Bpu1102I-digested pCRISPRia-v2 at 16 °C for 16 h. The ligation product was purified by isopropanol precipitation and then transformed into MegaX DH10B electrocompetent cells (Thermo Fisher Scientific) by electroporation using the Gene Pulser Xcell system (Bio-Rad), transforming ∼100 ng purified ligation product per 100 µL cells. The cells were allowed to recover in 3-6 mL SOC medium for 2 h. At that point, a small 1-5 uL aliquot was removed and plated in three serial dilutions on LB plates with selective antibiotic (carbenicillin). The remainder of the culture was inoculated into 0.5 to 1 L LB supplemented with 100 µg/mL carbenicillin, grown at 37 °C with shaking at 220 rpm for 16 h and harvested by centrifugation. Colonies on the plates were counted to confirm a transformation efficiency greater than 100-fold over the number of elements (>100x coverage). The pooled sgRNA plasmid library was extracted from the cells by GigaPrep (Qiagen or Zymo Research). Even coverage of library elements was confirmed by sequencing a small aliquot on a HiSeq 4000 (Illumina).

### Large-scale mismatched sgRNA screen and sequencing library preparation

Large-scale screens were conducted similarly to previously described screens^15, 19, 20^. The large-scale library was transduced in duplicate into K562 CRISPRi and Jurkat CRISPRi cells at MOI <1 (percentage of transduced cells 2 days after transduction: 20-40%) by centrifugation at 1000 × *g* and 33 °C for 2 h. Replicates were maintained separately in 0.5 L to 1 L of RPMI-1640 in 1 L spinner flasks for the course of the screen. 2 days after transduction, the cells were selected with puromycin for 2 days (K562: 2 days of 1 µg/mL; Jurkat: 1 day of 1 µg/mL and 1 day of 0.5 µg/mL), at which point transduced cells accounted for 80-95% of the population, as measured by flow cytometry using an LSR-II flow cytometer (BD Biosciences). Cells were allowed to recover for 1 day in the absence of puromycin. At this point, t_0_ samples with a 3000x library coverage (400 × 10^6^ cells) were harvested and the remaining cells were cultured further. The cells were maintained in spinner flasks by daily dilution to 0.5 × 10^6^ cells mL^−1^ at an average coverage of greater than 2000 cells per sgRNA with daily measurements of cell numbers and viability on an Accuri bench-top flow cytometer (BD BioSciences) for 11 days, at which point endpoint samples were harvested by centrifugation with 3000x library coverage.

Genomic DNA was isolated from frozen cell samples and the sgRNA-encoding region was enriched, amplified, and processed for sequencing essentially as described previously^19^. Briefly, genomic DNA was isolated using a NucleoSpin Blood XL kit (Macherey-Nagel), using 1 column per 100 × 10^6^ cells. The isolated genomic DNA was digested with 400 U SbfI-HF (NEB) per mg DNA at 37 °C for 16 h. To isolate the ∼500 bp fragment containing the sgRNA expression casette liberated by this digest, size separation was performed using large-scale gel electrophoresis with 0.8% agarose gels. The region containing DNA between 200 and 800 bp of size was excised and DNA was purified using the NucleoSpin Gel and PCR Clean-up kit (Macherey-Nagel). The isolated DNA was quantified using a QuBit Fluorometer (Thermo Fisher Scientific) and then amplified by 23 cycles of PCR using Phusion polymerase (NEB) and appending Illumina adapter and unique sample indices in the process. Each DNA sample was divided into 5-50 individual 100 µL reactions, each with 500 ng DNA as input. To ensure base diversity during sequencing, the samples were divided into two sets, with all samples for a given replicate always being assigned to the same set. The two sets had the Illumina adapters appended in opposite orientations, such that samples in set A were sequenced from the 5′ end of the sgRNA sequence in the first 20 cycles of sequencing and samples in set B were sequenced from the 3′ end of the sgRNA sequence in the next 20 cycles of sequencing. With updates to Illumina chemistry and software, this strategy is no longer required to ensure high sequencing quality, and all samples are amplified in the same orientation. Following the PCR, all reactions for a given DNA sample were combined and a small aliquot (100-300 µL) was purified using AMPure XP beads (Beckman-Coulter) with a two-sided selection (0.65x followed by 1x). Sequencing libraries from all samples were combined and sequencing was performed on a HiSeq 4000 (Illumina) using single-read 50 runs and with two custom sequencing primers (oCRISPRi_seq_V5 and oCRISPRi_seq_V4_3’, Table S15). For samples that were amplified in the same orientation, only a single custom sequencing primer was added (oCRISPRi_seq_V5), and the samples were supplemented with a 5% PhiX spike-in.

Sequencing reads were aligned to the library sequences, counted, and quantified using the Python-based ScreenProcessing pipeline (https://github.com/mhorlbeck/ScreenProcessing). Calculation of phenotypes was performed as described previously^15, 19, 20^. Untreated growth phenotypes (γ) were derived by calculating the log_2_ change in enrichment of an sgRNA in the endpoint and t_0_ samples, subtracting the equivalent median value for all non-targeting sgRNAs, and dividing by the number of doublings of the population^15, 20^. For sgRNAs with a read count of 0, a pseudocount of 1 was added. sgRNAs with <50 reads in both the endpoint and t_0_ samples in a given replicate were excluded from analysis. Read counts and phenotypes for individual sgRNAs are available in Table S3 and Table S4, respectively. To calculate relative activities, phenotypes of mismatched sgRNAs were divided by those for the corresponding perfectly matched sgRNA. Relative activities were filtered for series in which the perfectly matched sgRNA had a growth phenotype greater than 5 z-scores outside the distribution of negative control sgRNAs for all further analysis (3,147 and 2,029 sgRNA series for K562 and Jurkat cells, respectively). Relative activities from both cell lines were averaged if the series passed the z-score filter in both. All analyses were performed in Python 2.7 using a combination of Numpy (v1.14.0), Pandas (v0.23.4), and Scipy (v1.1.0).

### Design and pooled cloning of constant region variants library

The sequences in the library of modified constant regions were derived from the sgRNA (F+E) optimized sequence^23^ modified to include a BlpI site^15^. Each modified constant region was paired with 36 sgRNA targeting sequences (3 sgRNAs targeting each of 10 essential genes and six non-targeting negative control sgRNAs). The cloning strategy (described below) allowed the mutation of most positions in the sgRNA constant region. A variety of modifications were made, including substitutions of all single bases not in the BlpI restriction site (which is used for cloning), double substitutions including all substitutions at base-paired position pairs not before or in the BlpI site, and a variety of triple, quadruple, and sextuple substitutions, including base-pair-preserving substitutions at adjacent base-pairs.

The library was ordered and cloned in two parts. One part consisted of ∼100 modifications to the eight bases upstream of the BlpI restriction site. Constant region variants with mutations in this section were paired with each of the 36 targeting sequences, ordered as a pooled oligonucleotide library (Twist Biosciences), and cloned into pCRISPRia-v2 as described above. The second part consisted of ∼900 modifications to the 71 bases downstream of the BlpI restriction site. This part was cloned in two steps. First, all 36 targeting sequences were individually cloned into pCRISPRia-v2 as described above. The vectors were then pooled at an equimolar ratio and digested with BlpI (NEB) and XhoI (NEB). The modified constant region variants were ordered as a pooled oligonucleotide library (Twist Biosciences), PCR amplified with Phusion polymerase (NEB), digested with BlpI (NEB) and XhoI (NEB), and ligated into the digested vector pool, in a manner identical to previously published protocols and as described above, except for the different restriction enzymes.

### Compact mismatched sgRNA library and constant region library screens

Screens with the compact mismatched sgRNA library and the constant region library were conducted largely as described above, with smaller modifications during the screening procedure and an updated sequencing library preparation protocol. Briefly, the libraries were transduced in duplicate into K562 CRISPRi (both libraries) or HeLa CRISPRi cells (compact mismatched sgRNA library) as described above. K562 replicates were maintained separately in 0.15 to 0.3 L of RPMI-1640 in 0.3 L spinner flasks for the course of the screen. HeLa replicates were maintained in sets of ten 15-cm plates. Cells were selected with puromycin as described above (K562: 1 day of 0.75 µg/mL and 1 day of 0.85 µg/mL; HeLa: 2 days of 0.8 µg/mL and 1 day of 1 µg/mL). The remainder of the screen was carried out at >1000x library coverage (K562 compact mismatched sgRNA library: >2000x; HeLa compact mismatched sgRNA library: >1000x; K562 constant region library: >2000x). Multiple samples were harvested after 4 to 8 days of growth.

Genomic DNA was isolated from frozen cell samples as described above. The subsequent sequencing library preparation was simplified to omit the enrichment step by gel extraction. In particular, following the genomic DNA extraction, DNA was quantified by absorbance at 260 nm using a NanoDrop One spectrophotometer (Thermo Fisher Scientific) and then directly amplified by 22-23 cycles of PCR using NEBNext Ultra II Q5 PCR MasterMix (NEB), appending Illumina adapter and unique sample indices in the process. Each DNA sample was divided into 50-200 individual 100 µL reactions, each with 10 µg DNA as input. All samples were amplified using the same strategy and in the same orientation. The PCR products were purified as described above and sequencing libraries from all samples were combined. For the compact mismatched library screens, sequencing was performed on a HiSeq 4000 (Illumina) using single-read 50 runs with a 5% PhiX spike-in and a custom sequencing primer (oCRISPRi_seq_V5, Table S15). For the constant region screens, the PCR primers were adapted to allow for amplification of the entire constant region and to append a standard Illumina read 2 primer binding site (Table S15). Sequencing was then performed in the same manner including the custom sequencing primer (oCRISPRi_seq_v5) and a 5% PhiX spike-in, but using paired-read 150 runs.

Sequencing reads were processed as described above, except that sgRNAs with <50 reads (compact mismatched sgRNA library) or <25 reads (constant region library) in both the endpoint and t_0_ samples in a given replicate or with a read count of 0 in either sample were excluded from analysis. Read counts and phenotypes for individual sgRNAs are available in Tables S6-S7 (constant region screen) and Tables S10-S11 (compact mismatched sgRNA library screen).

### Generation and evaluation of individual constant region variants by RT-qPCR

Constant region variants were evaluated in the background of a constant region with an additional base pair substitution in the first stem loop (fourth base pair changed from AT to GC^25^). Ten constant region variants with average relative activities between 0.2 and 0.8 from the screen and carrying substitutions after the BlpI site were selected (Table S15). Cloning of individual constant regions was performed essentially as the cloning of sgRNA targeting regions, described above, except that the BlpI and XhoI restriction sites were used for cloning (the XhoI site is immediately downstream of the constant region) and that cloning was performed with a variant of pCRISPRia-v2 (marked with a puromycin resistance cassette and BFP, Addgene #84832)^19^. For each of the ten constant region variants as well as the constant region carrying only the stem loop substitution, two different targeting regions against *DPH2* were then cloned as described above (Table S1). These 22 vectors as well as a vector with a non-targeting negative control sgRNA (Table S1) were individually packaged into lentivirus and transduced into K562 CRISPRi cells at MOI < 1 (10 – 50% infected cells) by centrifugation at 1000 × *g* and 33 °C for 2 h. Cells were allowed to recover for 2 days and then selected to purity with puromycin (1.5 – 3 µg/mL), as assessed by measuring the fraction of BFP-positive cells by flow cytometry on an LSR-II (BD Biosciences), allowed to recover for 1 day, and harvested in aliquots of 0.5 – 2 × 10^6^ cells for RNA extraction. RNA was extracted using the RNeasy Mini kit (Qiagen) with on-column DNase digestion (Qiagen) and reverse-transcribed using SuperScript II Reverse Transcriptase (Thermo Fisher Scientific) with oligo(dT) primers in the presence of RNaseOUT Recombinant Ribonuclease Inhibitor (Thermo Fisher Scientific). Quantitative PCR (qPCR) reactions were performed in 22 µL reactions by adding 20 µL master mix containing 1.1x Colorless GoTaq Reaction Buffer (Promega), 0.7 mM MgCl_2_, dNTPs (0.2 mM each), primers (0.75 µM each), and 0.1x SYBR Green with GoTaq DNA polymerase (Promega) to 2 µL cDNA or water. Reactions were run on a LightCycler 480 Instrument (Roche). For each cDNA sample, reactions were set up with qPCR primers against *DPH2* and *ACTB* (sequences listed in Table S15). Experiments were performed in technical triplicates.

### Machine learning

In order to establish a subset of highly active sgRNAs with which to train a machine learning model, we filtered for perfectly matched sgRNAs with a growth phenotype greater than 10 z-scores outside the distribution of negative control sgRNAs in the K562 and/or Jurkat pooled screens (K562 γ < –0.21; Jurkat γ < –0.35). All singly mismatched variants derived from sgRNAs passing the filter were then included, and relative activities were calculated as described previously, averaging the replicate measurements for each sgRNA. In cases where a perfectly matched sgRNA passed the filter in the K562 and Jurkat screen, the average relative activity across both cell types was calculated for each mismatched variant; otherwise the relative activities for only one cell type were considered. This filtering scheme resulted in 26,248 mismatched sgRNAs comprising 2,034 series targeting 1,292 genes, with approximately 40% of relative activity values averaged from K562 and Jurkat cells.

For each sgRNA, a set of features was defined based on the sequences of the genomic target and the mismatched sgRNA. First, the genomic sequence extending from 22 bases 5′ of the beginning of the PAM to 1 base 3’ of the end of the PAM (26 bases in all) is binarized into a 2D array of shape (4, 26), with 0s and 1s indicating the absence or presence of a particular nucleotide at each position, respectively. Next, a similar array is constructed representing the mismatch imparted by the sgRNA, with an additional potential mismatch at the 5′ terminus of the sgRNA (position –20), which invariably begins with G in our libraries due to the mU6 promoter. Thus, the mismatched sequence array is identical to the genomic sequence array except for 1 or 2 positions. Finally, the arrays are stacked into a 3D volume of shape (4, 26, 2), which serves as the feature set for a particular sgRNA.

The training set of sgRNAs was established by randomly selecting 80% of sgRNA series, with the remaining 20% set aside for model validation. A convolutional neural network (CNN) regression model was then designed using Keras (https://keras.io/) with a TensorFlow backend engine, consisting of two sequential convolution layers, a max pooling layer, a flattening layer, and finally a three-layer fully connected network terminating in a single neuron. Additional regularization was achieved by adding dropout layers after the pooling step and between each fully connected layer. To penalize the model for ignoring under-represented sgRNA classes (e.g. those with intermediate relative activity), training sgRNAs were binned according to relative activity, and sample weights inversely proportional to the population in each bin were assigned. Hyperparameters were optimized using a randomized grid search with 3-fold cross-validation with the training set as input. Parameters included the size, shape, stride, and number of convolution filters, the pooling strategy, the number of neurons and layers in the dense network, the extent of dropout applied at each regularization step, the activation functions in each layer, the loss function, and the model optimizer. Ultimately, 20 CNN models with identical starting parameters were individually trained for 8 epochs in batches of 32 sgRNAs. Performance was assessed by computing the average prediction of the 20-model ensemble for each validation sgRNA and comparing it to the measured value.

A linear regression model was trained on the same set of sgRNAs, albeit with modified features more suited for this approach. These features include the identities of bases in and around the PAM, whether the invariant G at the 5′ end of the sgRNA is base paired, the GC content of the sgRNA, the change in GC content due to the point mutation, the location of the protospacer relative to the annotated transcription start site, the identities of the 3 RNA bases on either side of the mismatch, and the location and type of each mismatch. All features were binarized except for GC and delta GC content. In total, each sgRNA was represented by a vector of 270 features, 228 of which describe the mismatch position and type (19 possible positions by 12 possible types). Prior to training, feature vectors were z-normalized to set the mean to 0 and variance to 1. Finally, an elastic net linear regression model was created using the scikit-learn Python package (https://scikit-learn.org), and key hyperparameters (alpha and L1 ratio) were optimized using a grid search with 3-fold cross validation during training.

### Design of compact library

Genes targeted by the compact allelic series library were required to have at least one perfectly matched sgRNA with a growth phenotype greater than 2 z-scores outside the distribution of negative control sgRNAs (γ < –0.04) in a single replicate of a K562 pooled screen (this work or Horlbeck et al.^19^). By this metric, 4,722 unique sgRNAs targeting 2,405 essential genes were included. Next, for each perfectly matched sgRNA, variants containing all 57 single mismatches in the targeting sequence (positions –19 to –1) were generated *in silico*, and sequences with off-target binding potential in the human genome were filtered out as described for the large-scale library. Remaining variant sgRNAs were whitelisted for potential selection in subsequent steps.

For each gene being targeted, if both of the perfectly matched sgRNAs imparted growth phenotypes greater than 3 z-scores outside the distribution of negative controls (γ < –0.06) in this work’s large-scale K562 screen, then one series of 4 variant sgRNAs was generated from each. Otherwise, one series of 8 variants was generated from the sgRNA with the stronger phenotype. Both perfectly matched sgRNAs were included regardless of their growth phenotype, for a total of 2 perfectly matched and 8 mismatched sgRNAs per gene.

In order to select mismatched sgRNAs, we first divided the relative activity space into 6 bins with edges at 0.1, 0.3, 0.5, 0.7, and 0.9. For each series, we attempted to select sgRNAs from each of the middle 4 bins (centers at 0.2, 0.4, 0.6, and 0.8 relative activity) as measured in this work’s K562 screen. If multiple sgRNAs were available in a particular bin, they were prioritized based on distance to the center of the bin and variance between replicate measurements. If no previously measured sgRNA was available in a given bin, then the CNN model was run on all whitelisted (novel) mismatched sgRNAs belonging to that series, and sgRNAs were selected based on predicted activity as needed. In total, the compact library was composed of 4,722 unique perfectly matched sgRNAs, 19,210 unique mismatched sgRNAs, and 1,202 non-targeting control sgRNAs. Approximately 68% of mismatched sgRNAs were evaluated in previous screens (72% single mismatches, 28% double mismatches), with the remaining 32% imputed from the CNN model (all single mismatches). Sequences of sgRNAs and descriptions of mismatches are listed in Table S9.

### Perturb-seq

The Perturb-seq experiment targeted 25 genes involved in a diverse range of essential functions (Table S12). For each target gene, the original sgRNAs and 4-5 mismatched sgRNAs covering the range from full relative activity to low relative activity were chosen from the large-scale screen. These 128 targeting sgRNAs as well as 10 non-targeting negative control sgRNAs (Table S1) were individually cloned into a modified variant of the CROP-seq vector^33, 34^ as described above, except into the different vector. Lentivirus was individually packaged for each of the 138 sgRNAs and was harvested and frozen in array. To determine viral titers, each virus was individually transduced into K562 CRISPRi cells by centrifugation at 1000 × *g* and 33 °C for 2 h, and the fraction of transduced cells was quantified as BFP^+^ cells using an LSR-II flow cytometer (BD Biosciences) 48 h after transduction.

To generate transduced cells for single-cell RNA-seq analysis, virus for all 138 sgRNAs was pooled immediately before transduction and then transduced into K562 CRISPRi cells by centrifugation at 1000 × *g* and 33 °C for 2 h. To achieve even representation at the intended time of single-cell analysis, the virus pooling was adjusted both for titer and expected growth-rate defects. 3 d after transduction, transduced (BFP^+^) cells were selected using FACS on a FACSAria2 (BD Biosciences) and then resuspended in conditioned media (RPMI formulated as described above except supplemented with 20% FBS and 20% supernatant of an exponentially growing K562 culture). 2 d after sorting, the cells were loaded onto three lanes of a Chromium Single Cell 3’ V2 chip (10x Genomics) at 1000 cells/µL and processed according to the manufacturer’s instructions.

The CROP-seq sgRNA barcode was PCR amplified from the final single cell RNA-seq libraries with a primer specific to the sgRNA expression cassette (oBA503, Table S15) and a standard P5 primer (Table S15), purified on a Blue Pippin 1.5% agarose cassette (Sage Science) with size selection range 436-534 bp, and pooled with the single cell RNA-seq libraries at a ratio of 1:100. The libraries were sequenced on a HiSeq 4000 according to the manufacturer’s instructions (10x Genomics).

To measure the growth rate defects conferred by each sgRNA for comparison with the transcriptional phenotypes, samples of ∼500,000 transduced cells were taken from the same transduced cell population used in the Perturb-seq experiment on days two, seven, and twelve after transduction. Genomic DNA was extracted using the Nucleospin Blood kit (Macherey-Nagel) and sgRNA amplicons were prepared as described previously and above^19^, albeit with no genomic DNA digestion or gel purification, and sequenced on HiSeq 4000 as described above for the other screens. Growth phenotypes were calculated by comparing normalized sgRNA abundances at day seven and twelve to those at day two, as described above. Read counts and growth phenotypes (γ and relative activity) for individual sgRNAs are available in γ Table S13 and Table S14, respectively. Relative sgRNA activities measured at day seven (five days of growth) were used to assign sgRNA activities in further analysis.

### Perturb-seq data analysis

i) Cell barcode and UMI calling, assignment of perturbations UMI count tables with UMI counts for all genes in each individual cell were calculated from the raw sequencing data using CellRanger 2.1.1 (10x Genomics) with default settings. Perturbation calling was performed as described previously^27^. Briefly, reads from the specifically amplified sgRNA barcode libraries were aligned to a list of expected sgRNA barcode sequences using bowtie (flags: -v3 -q -m1). Reads with common UMI and barcode identity were then collapsed to counts for each cell barcode, producing a list of possible perturbation identities contained by that cell. A proposed perturbation identity was identified as “confident” if it met thresholds derived by examining the distributions of reads and UMIs across all cells and candidate identities: (1) reads > 50, (2) UMIs > 3, and (3) coverage (reads/UMI) in the upper mode of the observed distribution across all candidate identities. As described previously^44^, perturbation identities were called for any cell barcode with greater than 2,000 UMIs to enable capture of cells with strong growth defects. Any cell barcode containing two or more confident identities was deemed a “multiplet”, and may arise from either multiple infection or simultaneous encapsulation of more than one cell in a droplet during single-cell RNA sequencing. Cell barcodes passing the 2,000 UMI threshold and bearing a single, unambiguous perturbation barcode were included in all subsequent analyses.
ii) Expression normalization Some portions of analysis use normalized expression data. We used a relative normalization procedure based on comparison to the gene expression observed in control cells bearing non-targeting sgRNAs, as described previously^27^:

1. Total UMI counts for each cell barcode are normalized to have the median number of UMIs observed in control cells.
2. For each gene *x*, expression across all cell barcodes is z-normalized with respect to the mean (*μ_x_*) and standard deviation (σ*_x_*) observed in control cells:

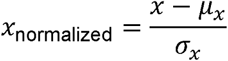 Following this normalization, control cells have average expression 0 (and standard deviation 1) for all genes. Negative/positive values therefore represent under/overexpression relative to control.
iii) Target gene quantification Expression levels of genes targeted by a given sgRNA were quantified by normalizing UMI counts of the targeted gene to the total UMI count for each individual cell (Fig. S8). Considering raw UMI counts of the targeted gene (Fig. S9) or z-normalized target gene expression as described above yielded similar results. Note that the sgRNA targeting *BCR* is toxic due to knockdown of the *BCR*-*ABL1* fusion present in K562 cells. Knockdown was apparent both in *BCR* and *ABL1* expression, but we used *BCR* expression for further analysis as there are likely additional copies of *ABL1* that are not fused to *BCR* (and thus would not be affected by the *BCR*-targeting sgRNA) contributing to *ABL1* expression.
iv) Cell cycle analysis Calling of cell cycle stages was performed using a similar approach to Macosko et al.^45^ and largely as described in Adamson and Norman et al.^27^. Briefly, lists of marker genes showing specific expression in different cell cycle stages from the literature were first adapted to K562 cells by restricting to those that showed highly correlated expression within our experiment. The total (log_2_-normalized) expression of each set of marker genes was used to create scores for each cell cycle stage within each cell, and these scores were then z-normalized across all cells. Each cell was assigned to the cell cycle stage with the highest score.
v) Differential gene expression analysis We took two approaches to differential expression, as described previously^44^. For both approaches, we only considered genes with expression greater than 0.25 UMIs per cell on average across all cells. First, for a given gene, we could assess the changes in the expression distribution of that gene induced by a given genetic perturbation by comparing to the expression distribution observed in control cells bearing non-targeting sgRNAs. We performed this comparison using a two-sample Kolmogorov-Smirnov test and corrected for multiple hypothesis testing at an FDR of 0.001 using the Benjamini-Yekutieli procedure. We also exploited a machine learning approach that potentially allows correlated expression patterns to be detected and that scales beyond two sample comparisons. Perturbed cells and control cells bearing non-targeting sgRNAs were each used as training data for a random forest classifier that was trained to predict which sgRNA a cell contained from its transcriptional state. As part of the training process the classifier ranks which genes have the most prognostic power in predicting sgRNA identity, which by construction will tend to vary across condition. For most further analysis, the top 100-300 genes by prognostic power were then considered.
vi) Constructing mean expression profiles For some analyses, expression profiles were averaged across all cells with the same perturbation. In general, this was done simply by calculating the mean z-normalized expression of all genes with mean expression level of 0.25 UMI or higher across all cells in the experiment or within the specific considered subpopulation (usually all cells with sgRNAs targeting a given gene as well as all control cells with non-targeting sgRNAs).
vii) UMAP Dimensionality reduction For UMAP dimensionality reduction^38^ of all cells, the 300 genes with the highest prognostic power in distinguishing cells by targeted gene as ranked by a random forest classifier were selected. Dimensionality reduction was then performed on the z-normalized single-cell expression profiles of these 300 genes using the following parameters: n_neighbors = 40, min_dist = 0.1, metric = ‘euclidean’, spread = 1.0. UMAP dimensionality reduction of subpopulations containing only cells with perturbation of a given gene or control cells was performed analogously but using the expression profiles of the 100 genes with the highest prognostic power and using n_neighbors = 15. From the UMAP projection, we concluded that ∼5% cells had misassigned sgRNA identities, as evident for example by the presence of cells with negative control sgRNAs within the cluster of cells with *HSPA5* knockdown. These cells had confidently assigned single perturbations and only expressed the corresponding barcode transcript, suggesting that they did not evade our doublet detection algorithm. We speculate that these cells expressed two different sgRNAs but silenced expression of one of the reporter transcripts. Given the strong trends in the results above, we concluded that this rate of misassignment did not substantially affect our ability to identify trends within cell populations.
viii) ISR scores Magnitude of ISR activation in individual cells was quantified as activation of the PERK (*EIF2AK3*) regulon from the gene set and activation coefficients determined previously^27^.

## Data Availability Statement

Raw and processed Perturb-seq data are available at GEO under accession code GSE132080. Raw and processed sgRNA read counts from pooled screens are provided as supplemental tables. All other data will be made available by the corresponding author upon request.

## Code Availability Statement

Custom scripts in this manuscript largely build on scripts published previously^19, 27, 44^. All custom scripts will be made available upon request.

## Acknowledgements

We thank Gregory Ow and Eric Collisson (UCSF) for sharing the mCherry-marked sgRNA expression vector, Ryan Pak, Jacob Stern, and Albert Xu for help with library cloning and sequencing library preparation, Britt Adamson for sharing the modified CROP-seq vector, Matt Jones, Jin Chen, Luke Gilbert, Joseph Replogle, and all members of the Weissman lab for helpful discussions, and Eric Chow, Derek Bogdanoff, and Kaitlin Chaung from the UCSF Center for Advanced Technology for help with sequencing. This work was funded by National Institutes of Health grants F32 GM116331 (MJ), P50 GM102706, U01 CA168370, R01 DA036858, and RM1 HG009490 (JSW), and R35 GM118061 (CAG) and the Innovative Genomics Institute, UC Berkeley (CAG). JSW is a Howard Hughes Medical Institute Investigator. DAS is supported by NSF Graduate Research Fellowship 1650113 and a Moritz-Heyman Discovery Fellowship. RAS is supported by a Fannie and John Hertz Foundation Fellowship and an NSF Graduate Research Fellowship. MAH is a Byers Family Discovery Fellow and is supported by the UCSF Medical Scientist Training Program and the School of Medicine. TMN is a fellow and JAH is the Rebecca Ridley Kry Fellow of the Damon Runyon Cancer Research Foundation (TMN: DRG-2211-15, JAH: DRG-2262-16).

## Author contributions

MJ conducted the large-scale growth screen, supervised the constant region and Perturb-seq experiments, implemented the linear machine learning model, analyzed data, conceived experiments, and wrote the manuscript. DAS conducted the GFP and constant region screens, implemented the deep learning model, designed and conducted the compact library screen, analyzed data, conceived experiments, and wrote the manuscript. RAS designed the constant region library and conducted a pilot screen, designed and conducted the Perturb-seq experiment, analyzed data, conceived experiments, and edited the manuscript. MAH assisted with the large-scale growth screen and with JSH designed the large-scale library. SMS evaluated modified constant region activities by RT-qPCR. JAH and TMN assisted with data analysis. CRL assisted with library cloning and screens. CAG supervised the generation of the large-scale library and edited the manuscript. JSW conceived and supervised experiments, and wrote the manuscript. All authors provided feedback on the manuscript.

## Supplemental Table Legends

**Table S1.** sgRNA sequences used in this study.

**Table S2.** Large-scale library sequences.

**Table S3.** Large-scale screen sgRNA counts.

**Table S4.** Large-scale screen phenotypes.

**Table S5.** Constant region library sequences.

**Table S6.** Constant region screen sgRNA counts.

**Table S7.** Constant region screen phenotypes.

**Table S8.** Genome-wide predicted sgRNA activities.

**Table S9.** Compact library sequences.

**Table S10.** Compact screen sgRNA counts.

**Table S11.** Compact screen phenotypes.

**Table S12.** Perturb-seq gene descriptions.

**Table S13.** Perturb-seq pooled growth sgRNA counts.

**Table S14.** Perturb-seq sgRNA sequences and pooled growth phenotypes (γ and relative activity).

**Table S15.** Oligonucleotide sequences used in this study.

## Supplemental figure legends

**Figure S1.**
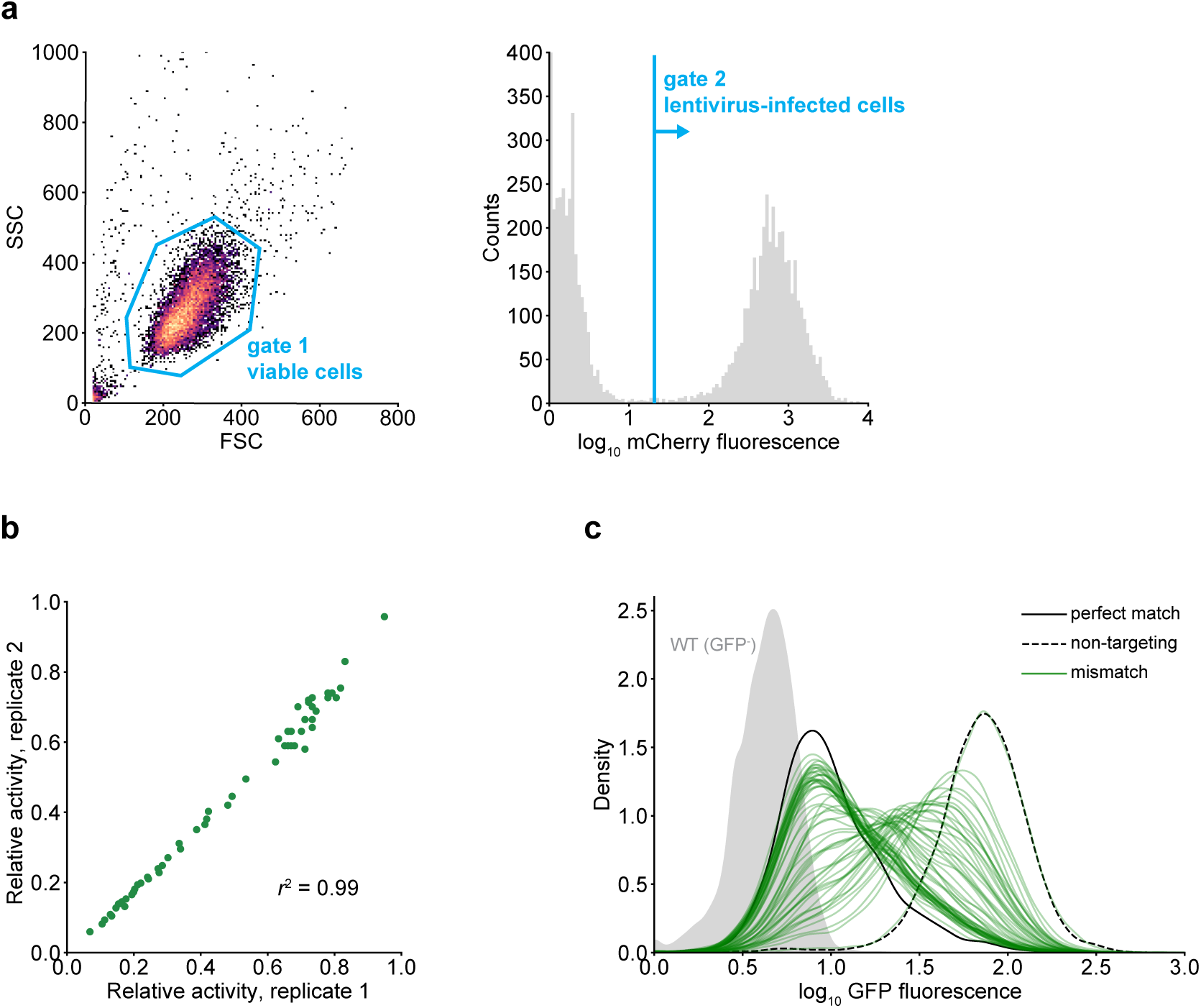
Details of the GFP mismatch experiment. **(a)** Representative plots illustrating gating strategy to select cells for analysis. **(b)** Comparison of relative activities measured in two biological replicates. Relative activity was defined as the fold-knockdown of each mismatched variant (GFP_sgRNA[non-targeting]_ / GFP_sgRNA[variant]_) divided by the fold-knockdown of the perfectly-matched sgRNA. The background fluorescence of a GFP^−^ strain was subtracted from all GFP values prior to other calculations. **(c)** KDE plots of GFP distributions 10 days after transducing K562 GFP^+^ cells with the perfectly-matched sgRNA, a non-targeting sgRNA, and each of the 57 singly-mismatched variants. Fluorescence of a GFP^−^ K562 strain is shown in gray. Although most GFP distributions are unimodal, some are broadened compared to those with the perfectly matched sgRNA or the negative control sgRNA. This heterogeneity could be a consequence of the random integration of the GFP locus, cell-to-cell differences in expression of the dCas9-KRAB effector in our polyclonal cell line, the amplification of gene expression bursts by long GFP half-lives, or a combination of these factors.

**Figure S2.**
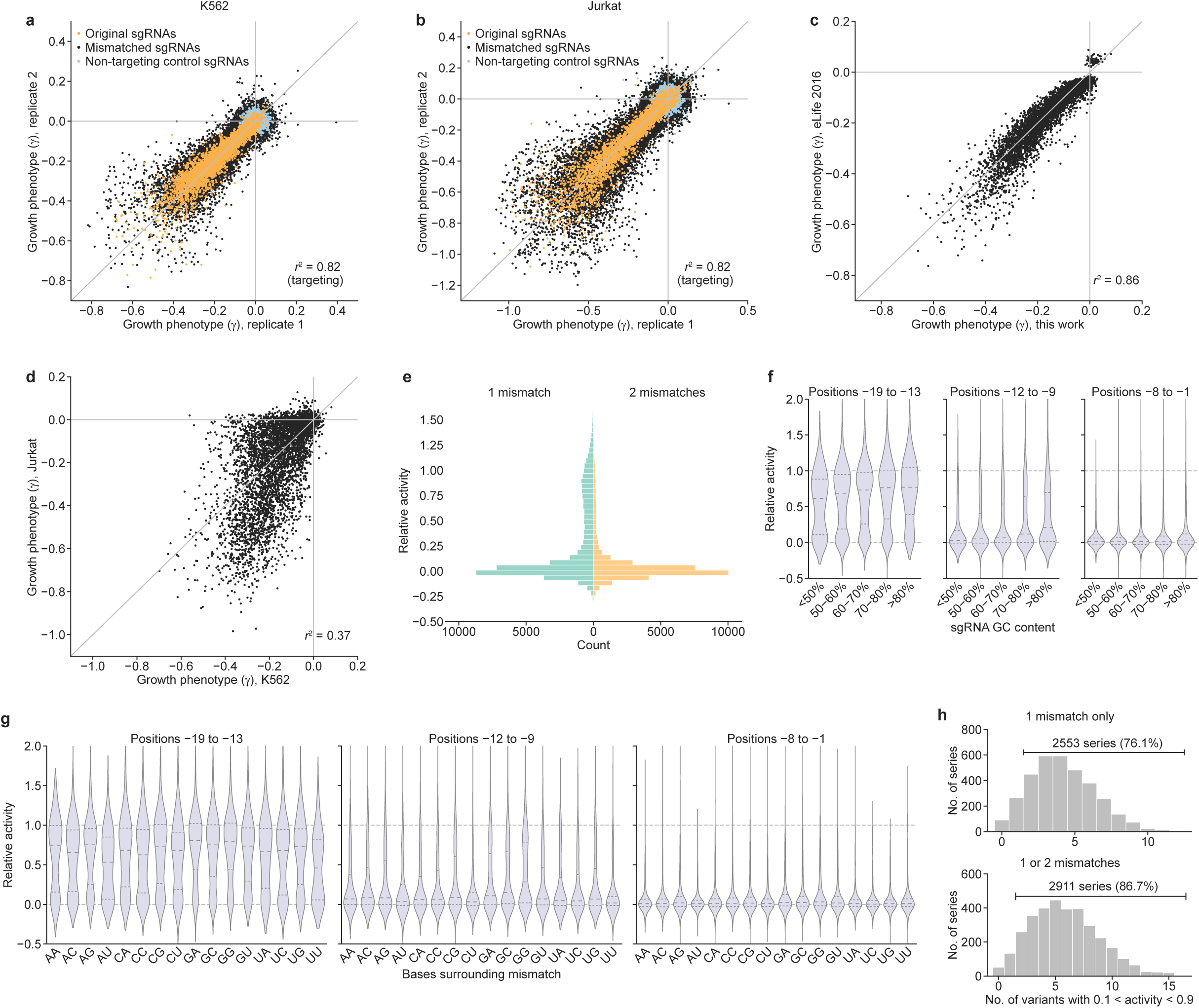
Additional analysis of large-scale mismatched sgRNA screen. **(a,b)** Comparison of growth phenotypes (γ of all sgRNAs derived from biological replicates of the **(a)** K562 and **(b)** Jurkat screens. **(c)** Comparison of growth phenotypes (γ) of perfectly matched sgRNAs from the γ K562 screen in this work and a previously published K562 screen^19^ (average of two biological replicates). **(d)** Comparison of growth phenotypes (γ) of perfectly matched sgRNAs in K562 and Jurkat cells reveals substantial differences, likely reflecting cell-type specific gene essentiality (average of two biological replicates). **(e)** Distribution of mismatched sgRNA relative activities for sgRNAs with 1 mismatch (left) or 2 mismatches (right). **(f)** Distribution of mismatched sgRNA relative activities stratified by sgRNA GC content, grouped by mismatches located in positions – 19 to –13 (PAM-distal region), positions –12 to –9 (intermediate region), and positions –8 to –1 (PAM-proximal/seed region). **(g)** Distribution of mismatched sgRNA relative activities stratified by the identity of the 2 bases flanking the mismatch, grouped by mismatches located in positions –19 to –13 (PAM-distal region), positions –12 to –9 (intermediate region), and positions –8 to –1 (PAM-proximal/seed region). **(h)** Distribution of sgRNA series by number of sgRNAs with intermediate activity (0.1 < relative activity < 0.9), using only sgRNAs with a single mismatch (top) or all mismatched sgRNAs (bottom).

**Figure S3.**
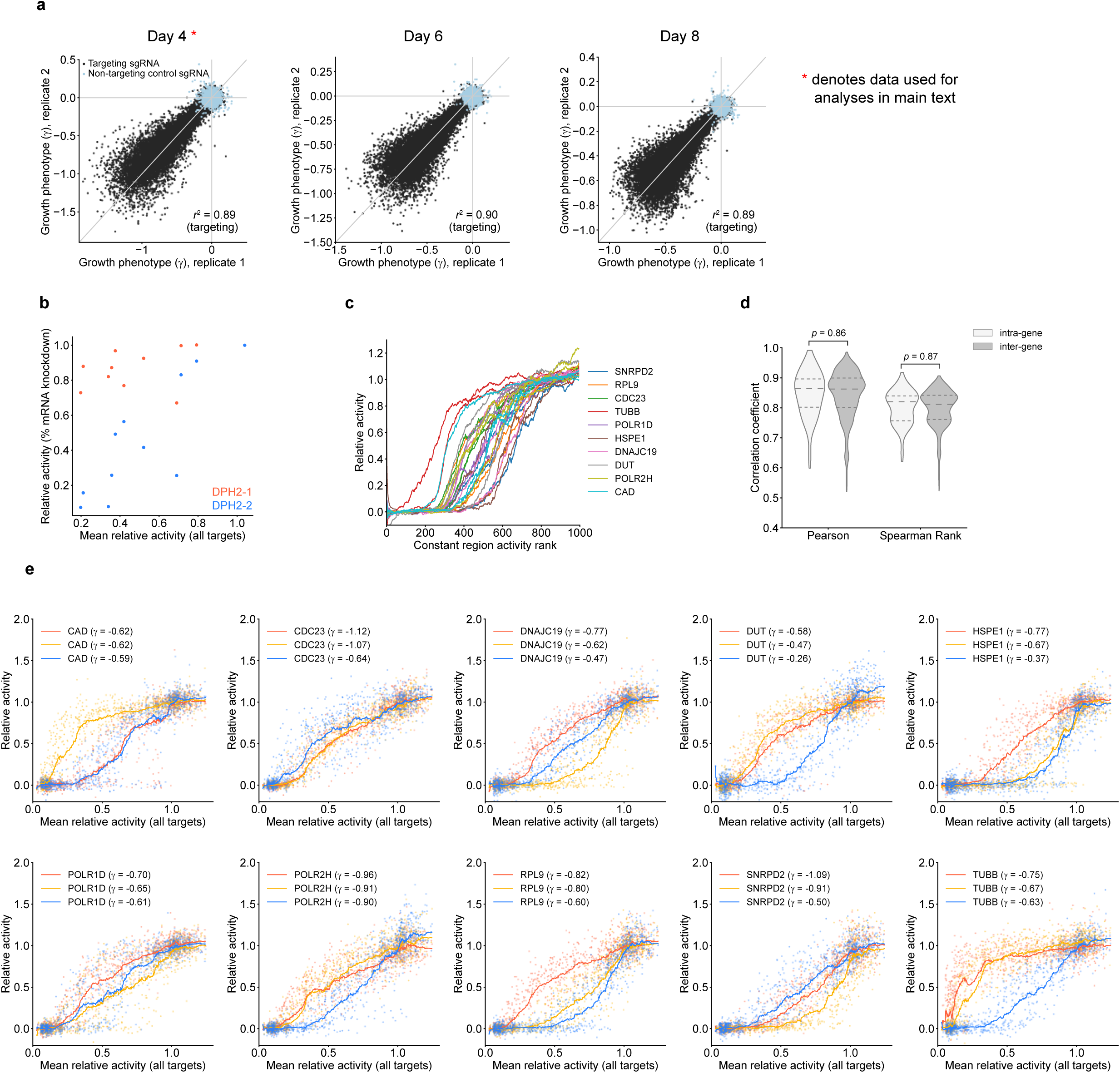
Additional analysis of modified constant regions. **(a)** Comparison of growth phenotypes measured in each biological replicate after 4, 6, or 8 days of growth from t_0_. Data from Day 4 was used for all subsequent analyses. **(b)** Comparison of relative % knockdown (quantified via RT-qPCR) and mean relative growth phenotype for 10 intermediate-activity constant region variants paired with two targeting sequences against *DPH2*. Data represent mean of technical triplicates. **(c)** Relative activities of constant regions paired with all 30 targeting sequences, ranked by the average strength of each constant region and displayed as rolling means with a window size of 50. **(d)** Distribution of all pairwise correlations of constant region relative activities within and between gene targets. Indicated *p*-values are derived from a two-tailed Student’s t-test. **(e)** Relative activity of each indicated target sequence:constant region pair vs. the mean relative activity of the respective constant region for all targets. Growth phenotypes (γ) with the unmodified constant region are indicated in the figure legends. Lines γ represent rolling means of individual data points.

**Figure S4.**
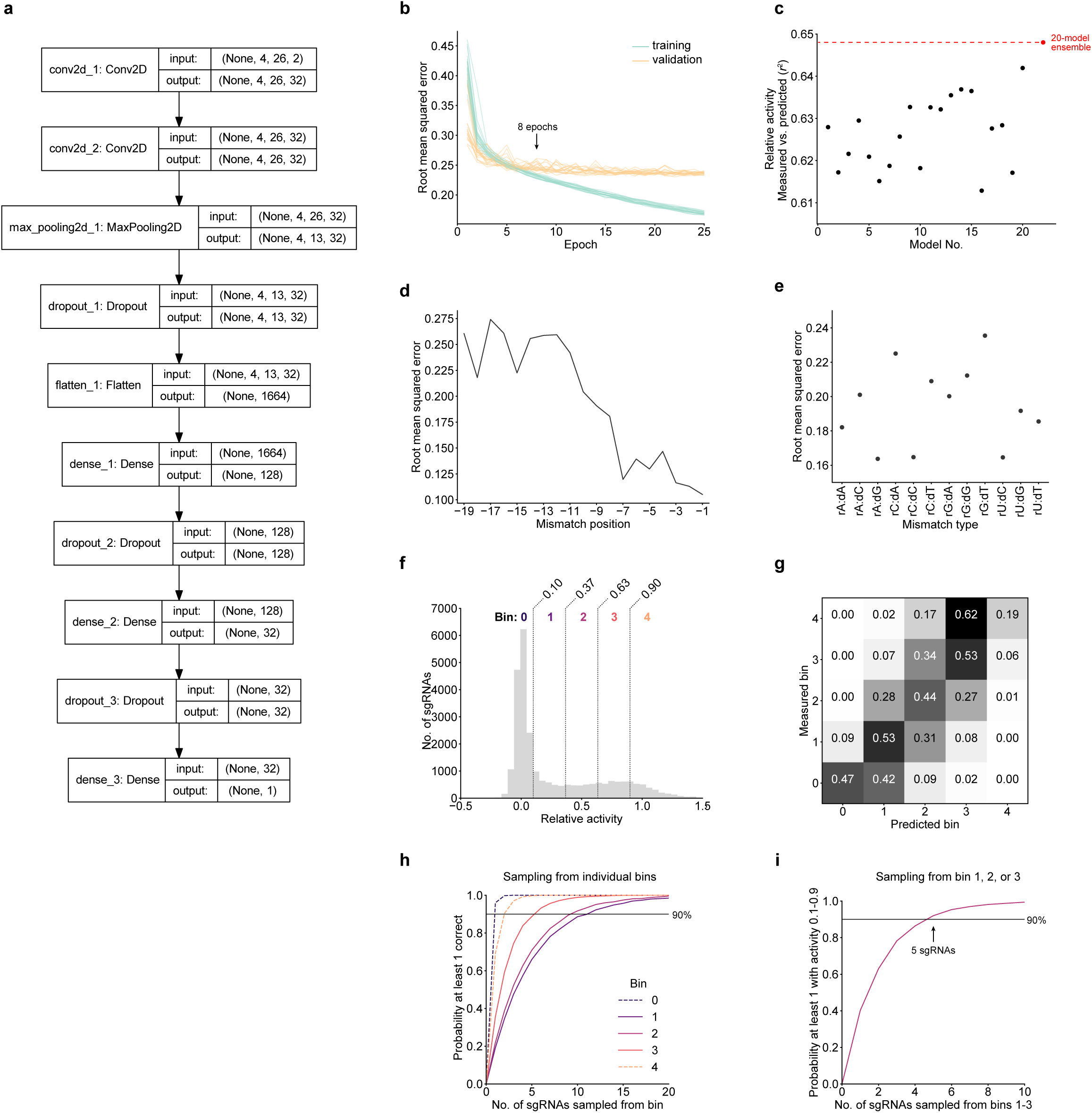
Additional details for the neural network. **(a)** Graph of the CNN model architecture. **(b)** Model loss, measured as root mean squared error, for training and validation data over 25 training epochs. Each line represents one of 20 models trained. The final models used for our predictions were only trained for 8 epochs, as additional cycles only reduced training loss without significant improvement in validation loss (i.e., the model becomes over-fit). **(c)** Explained variance (*r*^2^) of validation sgRNA relative activities for each individual model (black), and for the mean prediction of all 20 models (red). **(d)** Validation error stratified by mismatch position. **(e)** Validation error stratified by mismatch type. **(f)** Partitioning of sgRNAs into bins based on relative activity in the large-scale K562 screen. **(g)** Confusion matrix showing the fraction of sgRNAs in each actual (measured) activity bin that were assigned to each predicted bin by the CNN model. Each row sums to 1. **(h)** Statistics indicating the requisite number of randomly sampled sgRNAs from each activity bin to have a given probability of selecting at least one sgRNA with true activity in that bin. Simulations are based on the probabilities outlined in the confusion matrix (panel **e**). **(g)** Similar to panel **f**, with random sampling from any of the intermediate activity bins (1-3) to yield at least one sgRNA with intermediate activity (0.1-0.9).

**Figure S5.**
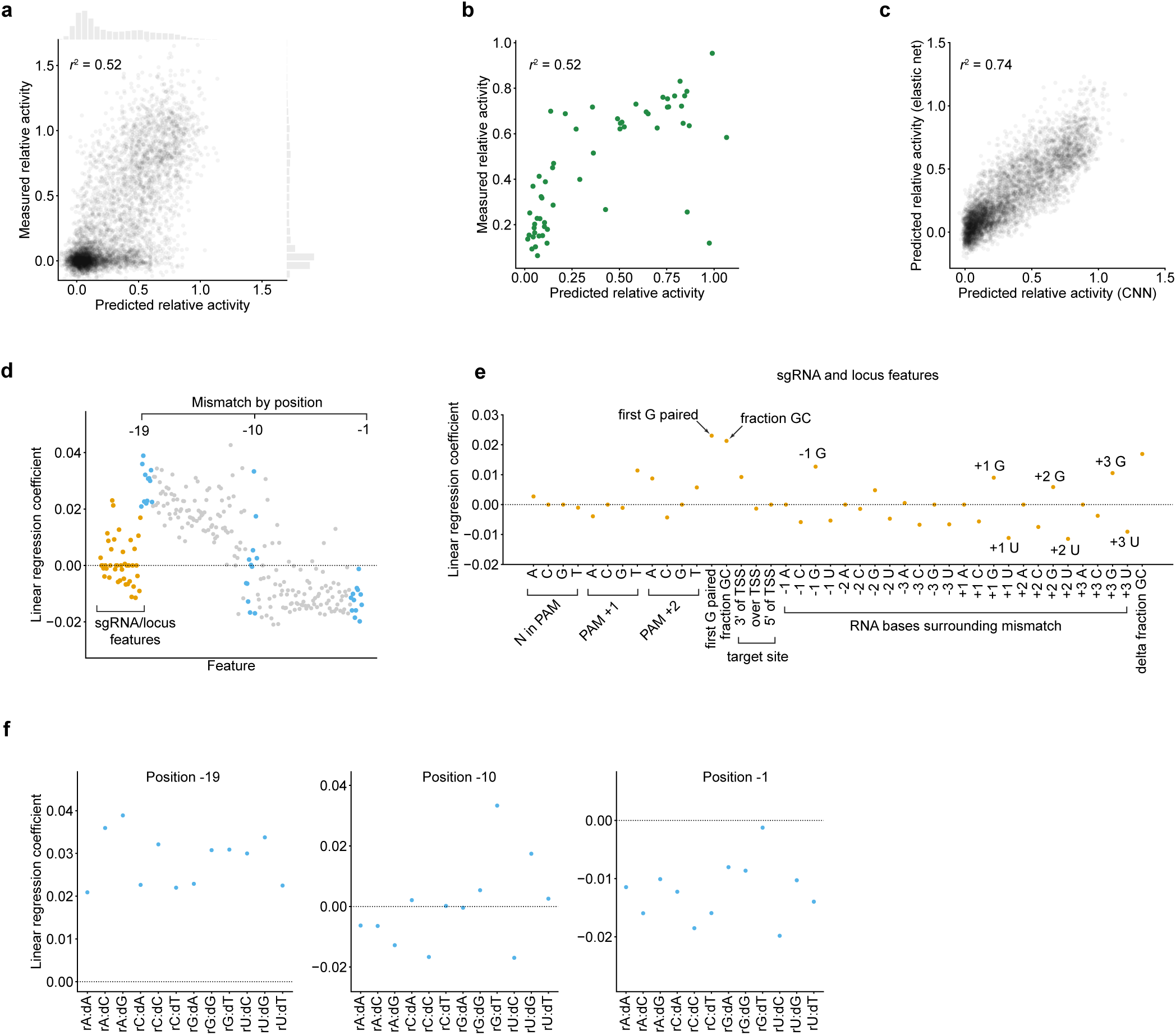
Additional details for the linear model. **(a)** Comparison of measured relative growth phenotypes from the large-scale screen and predicted activities assigned by the elastic net linear model. Marginal histograms show distributions of relative activities along the corresponding axes**. (b)** Comparison of measured relative activity (relative knockdown) in the GFP experiment and predicted relative sgRNA activity. **(c)** Comparison of predicted relative activities from the linear model and the neural network, based on the validation set of singly-mismatched sgRNAs. **(d)** Regression coefficients assigned to each feature in the linear model. 228 features (gray, blue) describe the position and type of mismatch; 42 features (gold) carry other information about the sgRNA and genomic context surrounding the protospacer. These features are detailed in subsequent panels. **(e)** Linear coefficients for features of the sgRNA and targeted locus. TSS; transcription start site. **(f)** Linear coefficients for features covering positions in the distal, intermediate, and seed regions of the targeting sequence (highlighted blue in panel **d**).

**Figure S6.**
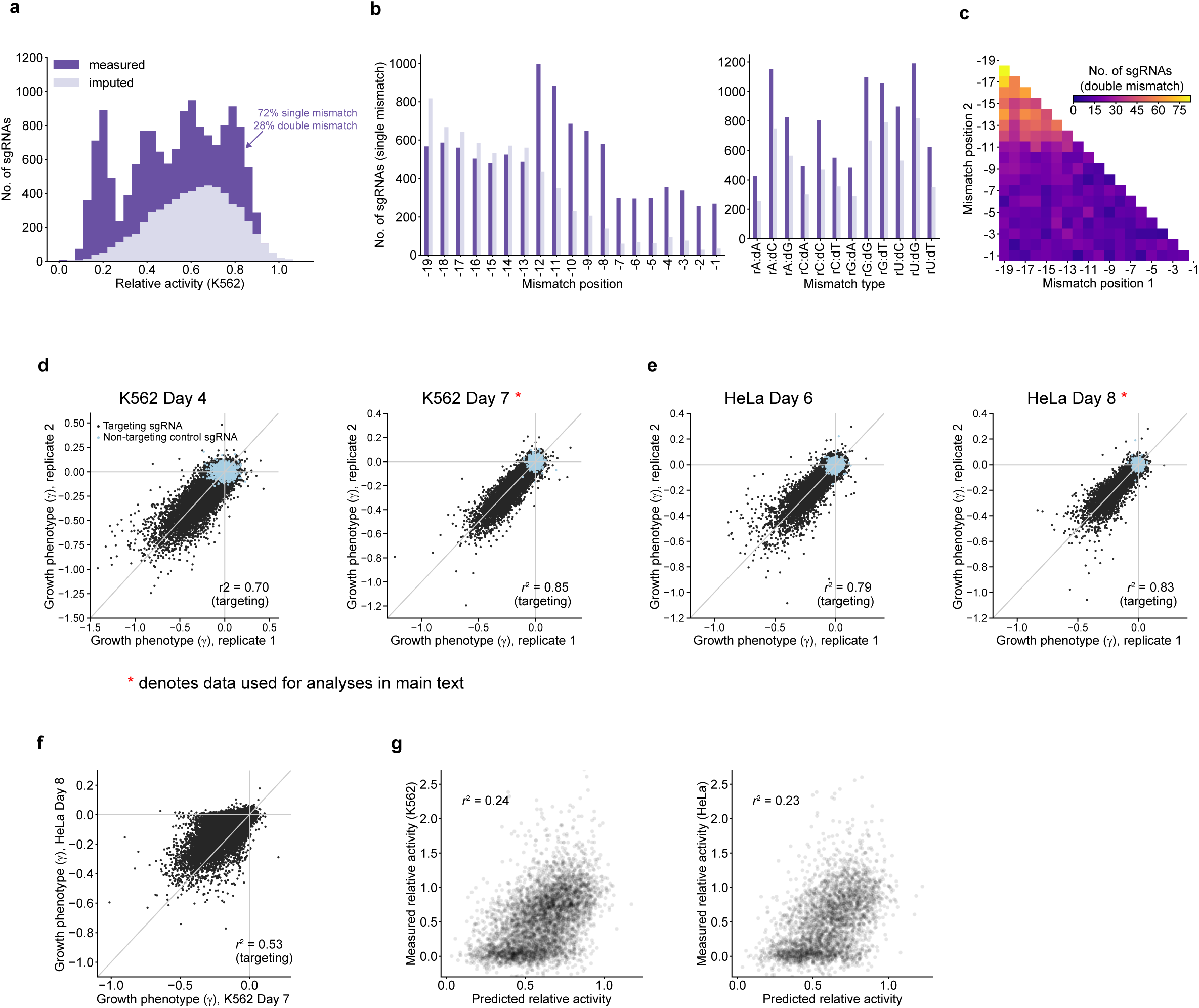
Additional analysis of the compact allelic series screen. **(a)** Composition of the compact library, in terms of previously measured relative activities in the large-scale screen (dark purple), or predicted relative activities assigned by the CNN model ensemble (light purple). Perfectly matched sgRNAs, which by definition have relative activities of 1.0, comprise 20% of the library but were not included in the histogram. **(b)** Distribution of mismatch positions and types for singly-mismatched sgRNAs in the compact library, for previously measured (dark purple) and CNN-imputed (light purple) sgRNAs. **(c)** Heatmap showing the distribution of mutated positions for doubly-mismatched sgRNAs in the compact library. **(d)** Comparison of growth phenotypes measured in each K562 biological replicate 4- and 7-days post-transduction. Data from Day 7 was used for all subsequent analyses. **(e)** Comparison of growth phenotypes measured in each HeLa biological replicate 6- and 8-days post-transduction. Data from Day 8 was used for all subsequent analyses. **(f)** Comparison of growth phenotypes in HeLa and K562 cells (γ expressed as the average of biological replicate measurements). **(g)** Measured vs. γ predicted relative activities of CNN-imputed sgRNAs in K562 cells (left) and HeLa cells (right). A small number of points beyond the y-axis limits were excluded to more clearly display the bulk of the distribution.

**Figure S7.**
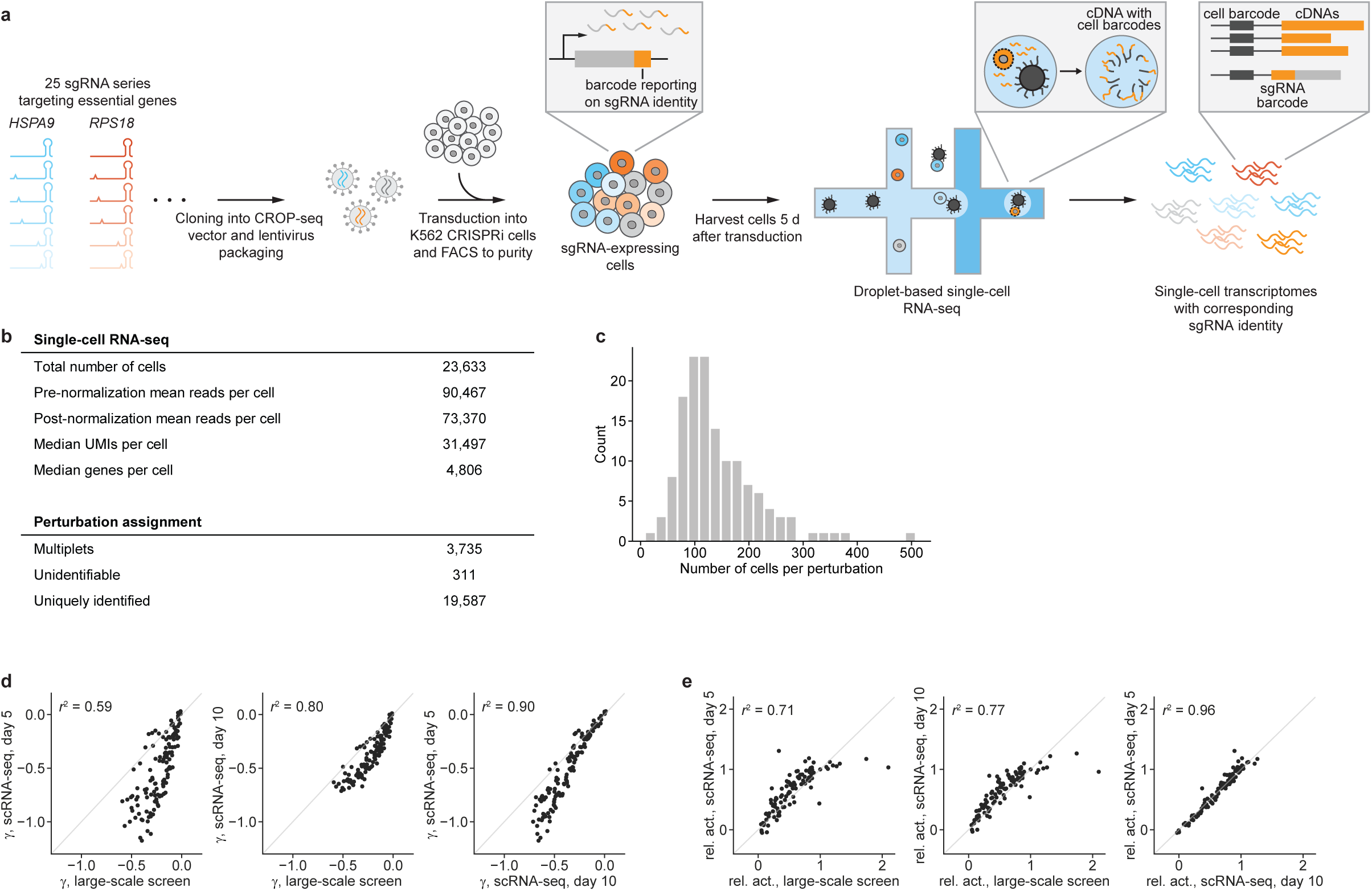
Summary of Perturb-seq experiment. **(a)** Schematic of Perturb-seq strategy to capture single-cell transcriptomes with matched sgRNA identities. **(b)** Summary of sequencing and perturbation assignment statistics. **(c)** Distribution of number of cells captured per perturbation. Median: 122 cells per perturbation; 5^th^ to 95^th^ percentile: 66 – 277 cells per perturbation. **(d,e)** Comparison of **(d)** growth phenotypes (γ) and **(e)** relative activities measured γ in the large-scale mismatched sgRNA screen and in the Perturb-seq experiment. Differences are likely due to the different timescales and the different vectors used.

**Figure S8.**
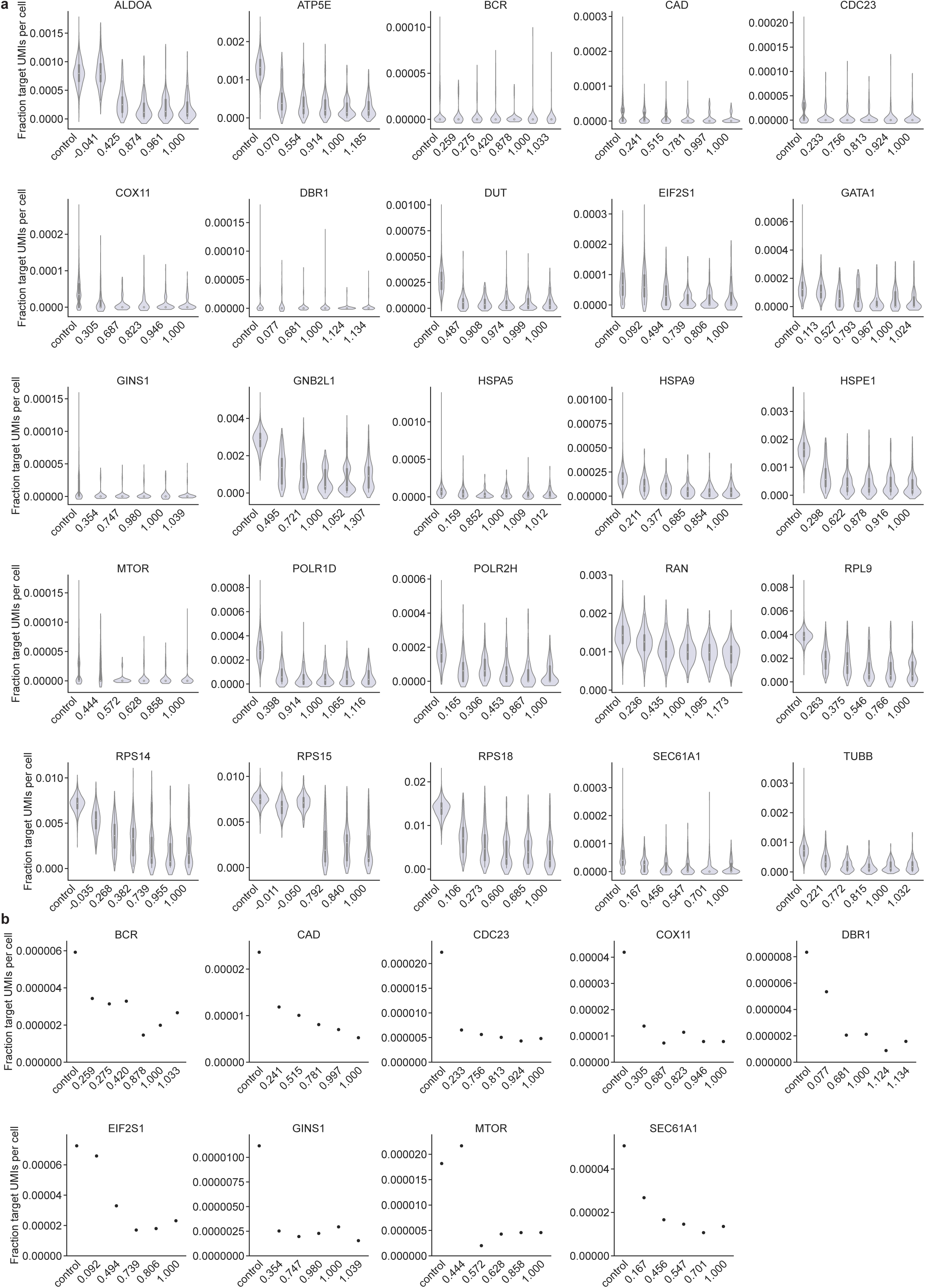
Target gene expression in cells with indicated perturbations. (**a**) Distribution of target gene expression levels, quantified as target gene UMI count normalized to total UMI count per cell. (**b**) Mean target gene expression levels for target genes with low basal expression levels.

**Figure S9.**
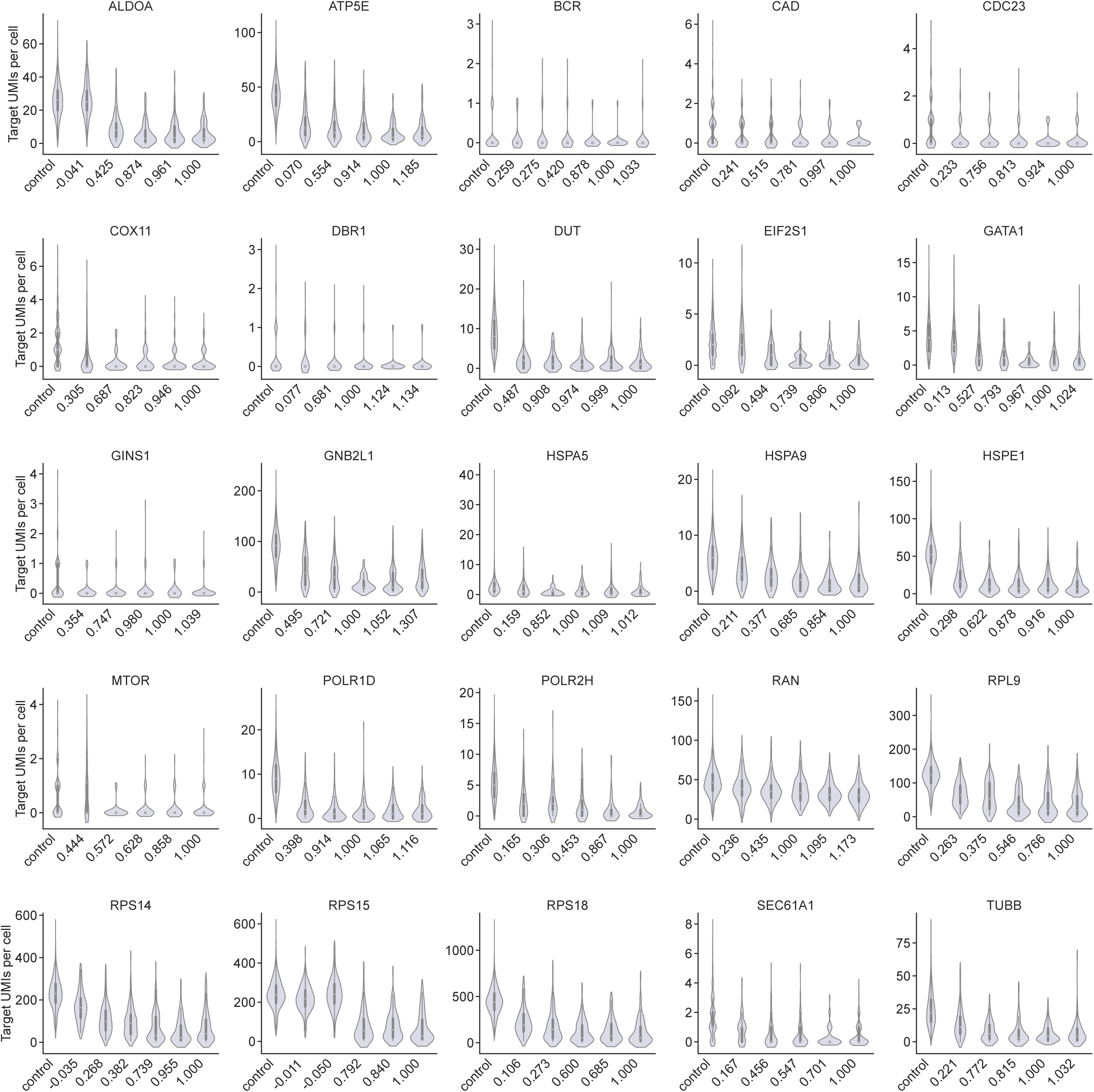
Distributions of target gene expression in cells with indicated perturbations. Expression is quantified as raw target gene UMI count.

**Figure S10.**
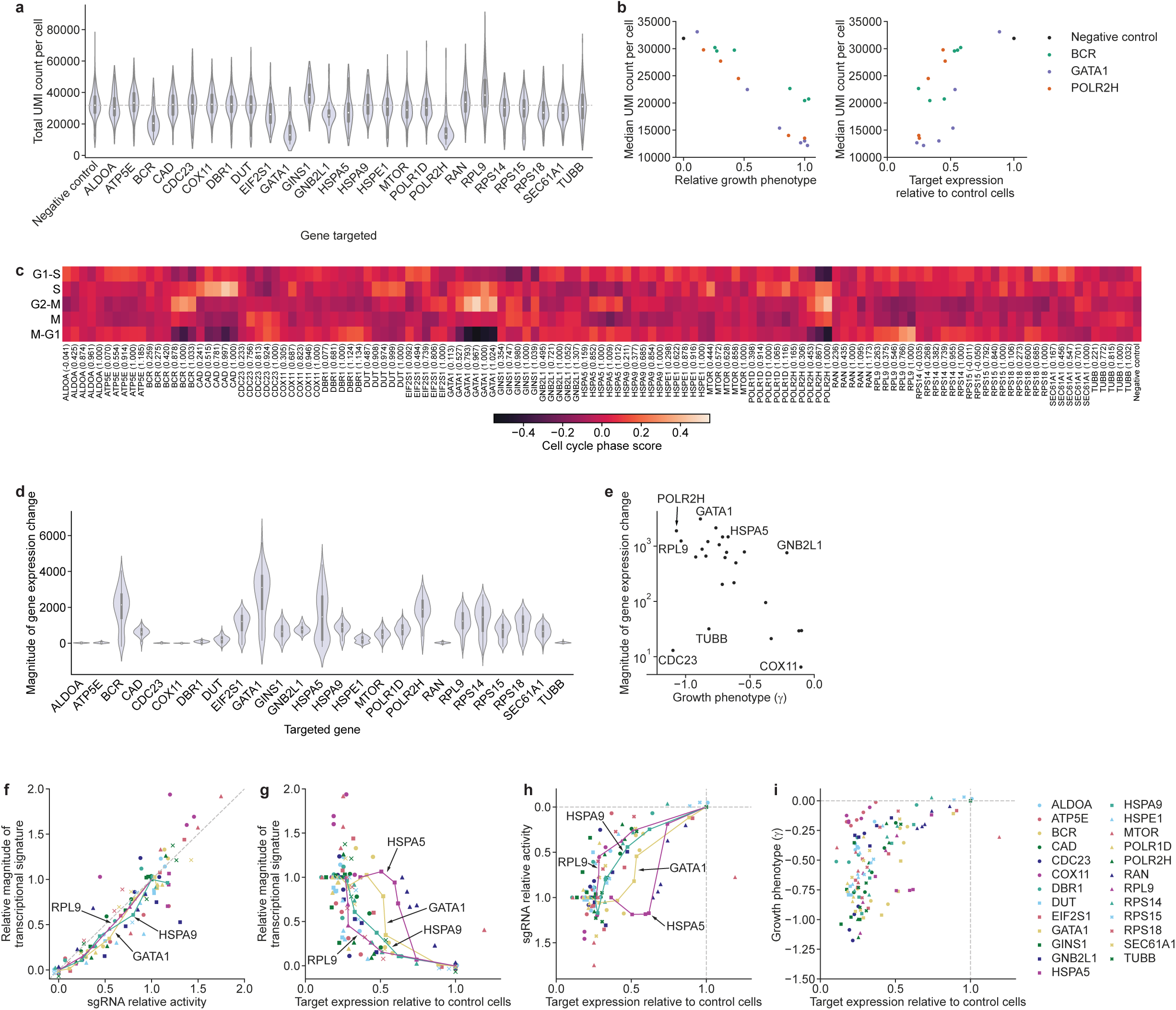
Phenotypes resulting from target gene titration. **(a)** Distributions of total UMI counts in cells with the perfectly matched sgRNA against the indicated genes. **(b)** Left: Comparison of median UMI count per cell and relative growth phenotype in cells with sgRNAs targeting *BCR*, *GATA1*, or *POLR2H* or control cells. Right: Comparison of median UMI count per cell and target gene expression. **(c)** Cell cycle scores (Methods) for populations of cells with individual sgRNAs. **(d)** Magnitudes of gene expression change of populations with perfectly matched sgRNAs targeting indicated genes. Magnitude of gene expression change is calculated as sum of z-scores of genes differentially expressed in the series (FDR-corrected *p* < 0.05 with any sgRNA in the series, two-sided Kolmogorov-Smirnov test, Methods), with z-scores of each gene in individual cells signed by the average direction of change in the population. **(e)** Comparison of magnitude of gene expression change to growth phenotype (γ) for all perfectly matched γ sgRNAs in the experiment. **(f)** Comparison of relative growth phenotype and magnitude of gene expression change for all individual sgRNAs, as in Fig. 6f but without increased transparency for individual series. **(g)** Comparison of magnitude of gene expression and target gene knockdown, as in Fig. 6g but without increased transparency for individual series. **(h)** Comparison of relative growth phenotype and target gene expression, as in Fig. 6f. **(i)** Comparison of measured growth phenotype (γ, not normalized to strongest sgRNA) and target gene expression, as in Fig. 6f.

**Figure S11.**
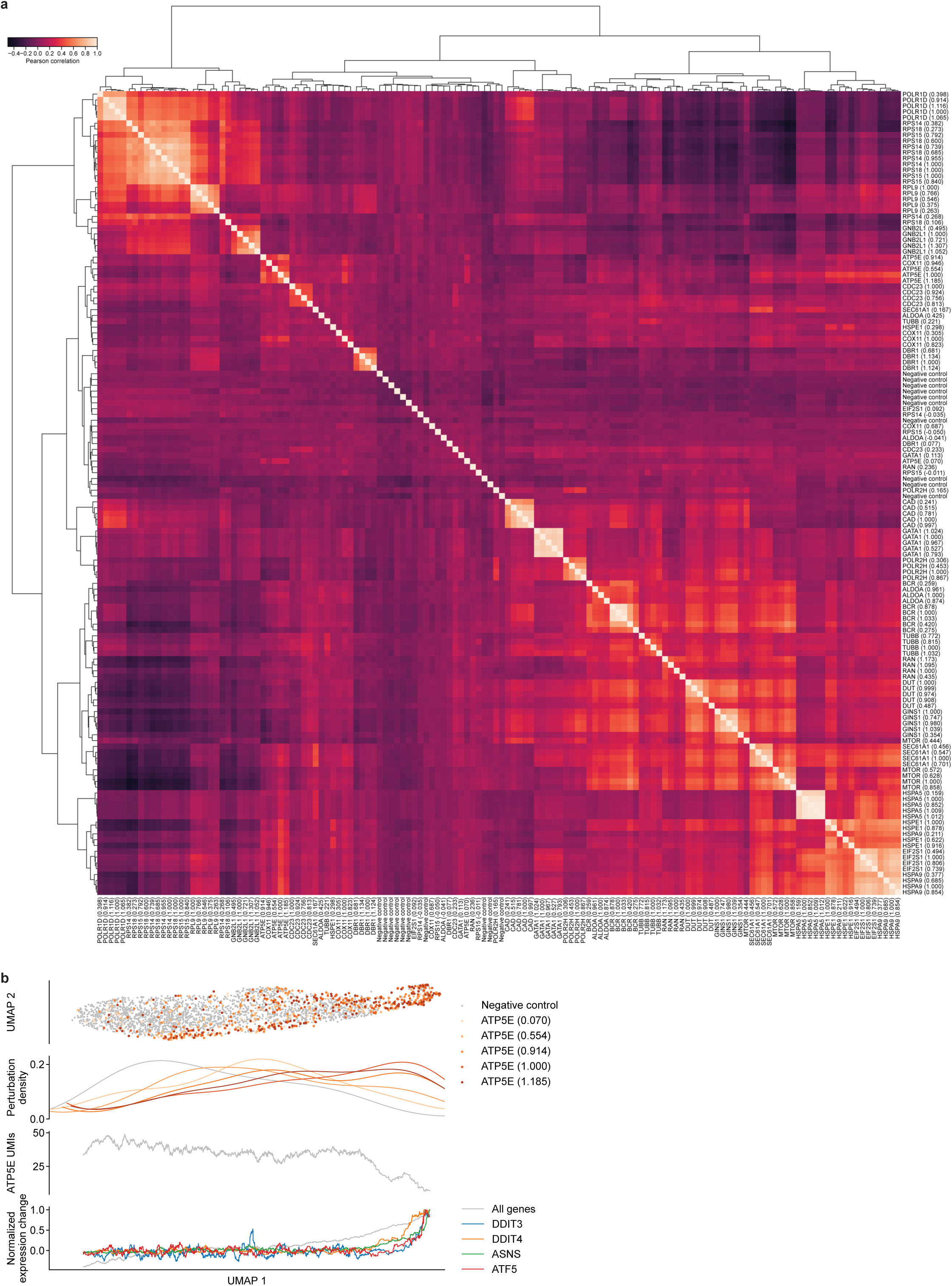
Diverse phenotypes resulting from essential gene depletion. (a) Clustered correlation heatmap of perturbations. Gene expression profiles for genes with mean UMI count > 0.25 in the entire population were z-normalized to expression values in cells with negative control sgRNAs and then averaged for populations with the same sgRNA. Crosswise Pearson correlations of all averaged transcriptomes were clustered by the Ward variance minimization algorithm implemented in scipy. **(b)** UMAP projection, distribution of cells with indicated sgRNAs, target gene expression (rolling mean over 50 cells), and magnitudes of transcriptional changes for all differentially expressed genes and selected ISR regulon genes (rolling mean over 50 cells) for cells with knockdown of *ATP5E* or control cells.

